# GPCR-mediated clearance of tau in post-synaptic compartments attenuates tau pathology *in vivo*

**DOI:** 10.1101/2020.01.21.914135

**Authors:** Ari W. Schaler, Avery M. Runyan, Stephanie L. Fowler, Helen Y. Figueroa, Seiji Shioda, Ismael Santa-Maria, Karen E. Duff, Natura Myeku

## Abstract

Accumulation of pathological tau in synapses has been identified as an early pathogenic event in Alzheimer’s disease (AD) and correlates strongly with cognitive decline in patients with AD. Tau is a cytosolic, axonal protein. However, in the disease condition, tau accumulates in post-synaptic compartments and pre-synaptic terminals, either due to missorting within neurons, trans-synaptic transfer between neurons or due to failure of clearance systems in synapses. Using a sub-cellular fractionation assay, we show that progressive deposition of seed competent tau occurs predominantly in post-synaptic compartments in a tau transgenic mouse and in AD patient brain, making these neuronal structures particularly vulnerable to tau toxicity. Tau-mediated post-synaptic toxicity could be further exacerbated by impaired proteasome activity which we detected by measuring the levels of polyubiquitin chains that target proteins to proteasomal degradation. To combat the accumulation of tau and proteasome impairment at the subcellular level, we devised a therapeutic strategy of proteasome-mediated clearance of tau restricted to the post-synaptic compartment. Utilizing the pharmacology of GPCRs, we show that *in vivo* stimulation of the PAC1R receptor by its ligand can propagate intracellular PKA signaling leading to enhanced synaptic proteasome activity and reduced tau in the post-synaptic compartment. Over time, clearance of post-synaptic tau led to reduced tauopathy and cognitive decline in rTg4510 mice. Together, these results highlight a novel therapeutic strategy of targeting GPCRs that propagate cAMP/PKA signaling as a tool to activate proteolysis restricted to synapses to prevent the accumulation of tau in the early stages of AD.

## Introduction

Synaptic dysfunction and synaptic spine loss is the strongest pathological correlate of cognitive decline in AD (*1–5*), with increasing data implicating the accumulation of synaptic tau in the progression of the disease (*6–9*). Recent evidence from AD post-mortem tissue suggests that the accumulation of tau oligomers in synapses correlated better with cognitive decline than the accumulation of Aβ oligomers, plaque burden, or somatic tau tangles (*10*). Although low levels of tau are present in pre-synaptic terminals and dendritic spines under physiological conditions (*11*), the accumulation of pathological tau in dendrites is a long-standing observation that has been identified as an early pathogenic event in AD (*12, 13*). How normal cytosolic tau that maintains higher concentration in axons is distributed to dendrites and synaptic spines in the disease condition is not well understood. However, studies with tauopathy animal models indicate that hyper-phosphorylation of tau can cause tau to mislocalize to dendrites and dendritic spines (*14–16*). Moreover, the manifestation of the cell-to-cell spread of tau species can further drive synaptic toxicity. Studies in live patients with AD and progressive supranuclear palsy (PSP) (*17*), and numerous *in vivo* animal studies (*18–27*) have recently demonstrated what was described first by Braak *et al*. that pathological tau species spread in a hierarchical pattern along the anatomical connection in AD (*28*).

The accumulation of pathological tau in synapses correlates with the accumulation of ubiquitinated proteins, suggesting disruption of the ubiquitin proteasome system (UPS) at synaptic compartments (*29*). The UPS machinery in synapses is critical for normal functioning of synapses, including synaptic protein turnover (*30–32*), plasticity (*33, 34*), and long-term memory formation (*35–37*), which rely on tightly controlled changes in the proteome. Recent studies in primary neuronal cultures using microfluidic chambers showed that the protein degradation systems (UPS and autophagy) play an essential role in the distribution of tau to dendrites and dendritic spines (*15*). The study found that local inhibition of protein degradation restricted to neurites led to missorting of phosphorylated tau to dendrites and loss of dendritic spines, whereas enhancing protein degradation pathways reduced tau missorting below basal levels (*15*). Degradation of tau and other poly-ubiquitinated substrates is carried out by the 26S proteasome, a 2.5 MDa, multi-catalytic, ATP-dependent protease that degrades proteins into small peptides. It is comprised of two components: a 19S regulatory particle (RP) and a 20S core particle (CP) that carries out the catalytic activity (*38*). In addition to the accumulation of ubiquitinated proteins, decreased proteasome activity has been reported in the hippocampal, parietal and temporal lobe regions of post-mortem AD brain tissue, but not in unaffected areas such as occipital lobe and cerebellum (*39*). The mechanism by which the proteasome becomes dysfunctional has been addressed by our previous studies (*40*) and by several other independent studies (*41–47*), suggesting that aggregates may disrupt protein degradation by physically blocking the gate opening of the 19S regulatory subunit of the 26S proteasome. A recent detailed mechanistic study showed that oligomers from different neurodegenerative diseases could impair proteasome function by binding to the outer surface of the 20S core particle and allosterically stabilizing the closed gate conformational state of the proteasome and blocking protein degradation (*47*). These data suggest that impairment of proteasome activity in various neurodegenerative diseases may be a common mechanism. Moreover, we have shown that 26S proteasomes remained defective in the degradation of ubiquitinated proteins even after they had been purified from the brains of the rTg4510 tauopathy mouse model that develops robust tau aggregates (*40*). Therefore, the evidence suggests that the persistence of proteasome deficiency in tauopathy caused by tau aggregates can cause fundamental changes in the quaternary structure of the 26S complex leading to a profound deterioration of its function.

Unless they are removed, toxic tau aggregates can disrupt the proper function not only of the UPS but of the nervous system. To prevent impairment of the proteasome-mediated degradation associated with tauopathy, we tested whether proteasome activity could be directly enhanced *in vivo* through cAMP/PKA-mediated phosphorylation, which we achieved by administration of PDE inhibitors (rolipram or cilostazol) (*40, 48*). Enhanced proteolysis resulted in attenuation of tauopathy and rescued cognitive decline in the rTg4510 mouse line. Other independent studies have confirmed that phosphorylation of proteasome subunits by protein kinase A (PKA) (*49, 50*) and other kinases such as protein kinase G (PKG) (*51, 52*) and dual-specificity tyrosine-regulated kinase 2 (DYRK2) (*53, 54*) can enhance proteasome-mediated protein degradation similarly. Therefore, the possibility of proteasome activity as a drug target is especially appealing as it can be applied to several proteinopathy diseases. The extent to which proteasome activation is required to maintain safety and therapeutic efficacy is not known. However, in general, cell-wide activation of protein degradation by proteasome or autophagy may not be a desirable strategy for long-term treatment of chronic neurodegenerative disease. Thus, targeting toxic tau in the specific neuronal compartments that are reported to accumulate tau in the early stage of AD before immunohistochemical detection of somatic NFT, could potentially be a safer strategy to prevent overt aggregation and spread of tau.

One therapeutic strategy that we devised for this study is the clearance of tau restricted to the post-synaptic compartment. By utilizing the pharmacology of G protein-coupled receptors (GPCRs) that mediate activation of adenylyl cyclase (AC), production of cyclic adenosine monophosphate (cAMP) and stimulation of PKA with region, cell type and sub-cellular -specific patterns (*55*), we hypothesized that stimulation of particular GPCRs can lead to PKA-dependent clearance of toxic tau species accumulated in dendritic spines and dendrites. We envisaged that this is a plausible hypothesis because GPCRs that mediate slow synaptic transmission by modulating the release and binding of neurotransmitters (*55–57*) are situated on the pre and post-synaptic terminals (*55*) and can propagate signal transduction restricted to synapses.

In our study, we tested whether stimulation of pituitary adenylate cyclase-activating polypeptide (PACAP) type 1 receptor (PAC1R), a Gs-coupled GPCR, present on the membrane of the post-synaptic compartment can stimulate cAMP/PKA/proteasome tau degradation predominantly in the post-synaptic compartment. In the brain, PAC1R is stimulated by its ligand PACAP, which acts as a neurotransmitter, neurotrophic factor, and a neurohormone (*58, 59*). Its effect on cell survival is mediated through stimulation of AC/cAMP/PKA signaling (*60*). In mammals, two biologically active forms of PACAP have been identified. The predominant form in CNS is PACAP38, composed of 38 amino acids, and the minor form is PACAP27, composed of the first 27 amino acids of PACAP38 (*59*).

In this study, the subcellular characterization of tau species across different stages of tauopathy in rTg4510 mice and post-mortem tissue from AD patients shows that post-synaptic compartments are highly vulnerable to tau pathology due to accumulation of seed competent, high molecular weight (HMW), AT8 positive tau forms. Moreover, the chronic stimulation of PAC1R by PACAP38 (hereafter PACAP) resulted in proteasome-mediated clearance of post-synaptic tau, overall reduced hyper-phosphorylated and aggregated tau and improved cognitive performance in rTg4510 mice.

## RESULTS

### Deposition of pathological tau in post-synaptic compartments

Physiological tau is present predominantly in the axon of neurons with low levels present in the soma, dendrites, and pre-synaptic axon terminals (*61–64*). In contrast, synaptosomes isolated from AD brains contain high levels of hyper-phosphorylated (*11, 65*) and ubiquitinated (*29*) insoluble tau forms. Studies on primary neuronal cultures (*15, 66, 67*) and tauopathy mouse models (*14, 68, 69*) support the mechanism of missorting of tau to dendrites/post-synaptic compartments in early stages of the disease leading to disruption of synaptic function (*70–72*), loss of dendritic spines (*14, 67, 73, 74*) and cognitive impairment (*14, 73, 75, 76*). It is thought that the deposition of tau to dendrites marks an important event in AD due to its ability to mediate amyloid-β toxicity inside neurons (*74*). Despite mounting evidence that the accumulation of abnormal tau is synaptotoxic, it is not clear how tau pathology leads to synaptic dysfunction and loss.

To study the spatial and temporal distribution of tau species across tauopathy stages in rTg4510 mice, we used a method of sub-cellular fractionation in sucrose isotonic buffers to isolate synaptosomes and sub-synaptic compartments from a cytosolic fraction (**Supplementary Fig. 1A** (schematic diagram)). These structures form instantaneously when tissue is homogenized in sucrose isotonic buffers, whereby synaptic boutons reseal to form spherical structures known as synaptosomes. To investigate the distribution of tau within synapses, gradient-purified synaptosomes were further separated into pre and post -synaptic compartments. Thus, from the total extracts we generated cytosolic, synaptic, pre and post -synaptic fractions from WT (3 and 8 months of age) and rTg4510 mice across three stages of tauopathy: early (3 months), mid (5 months) and late-stage (8 months) from intact cortical brain tissue (**Fig. 1A**, for uncut blots, see **Supplementary Fig. 1B**).

**Figure 1.**
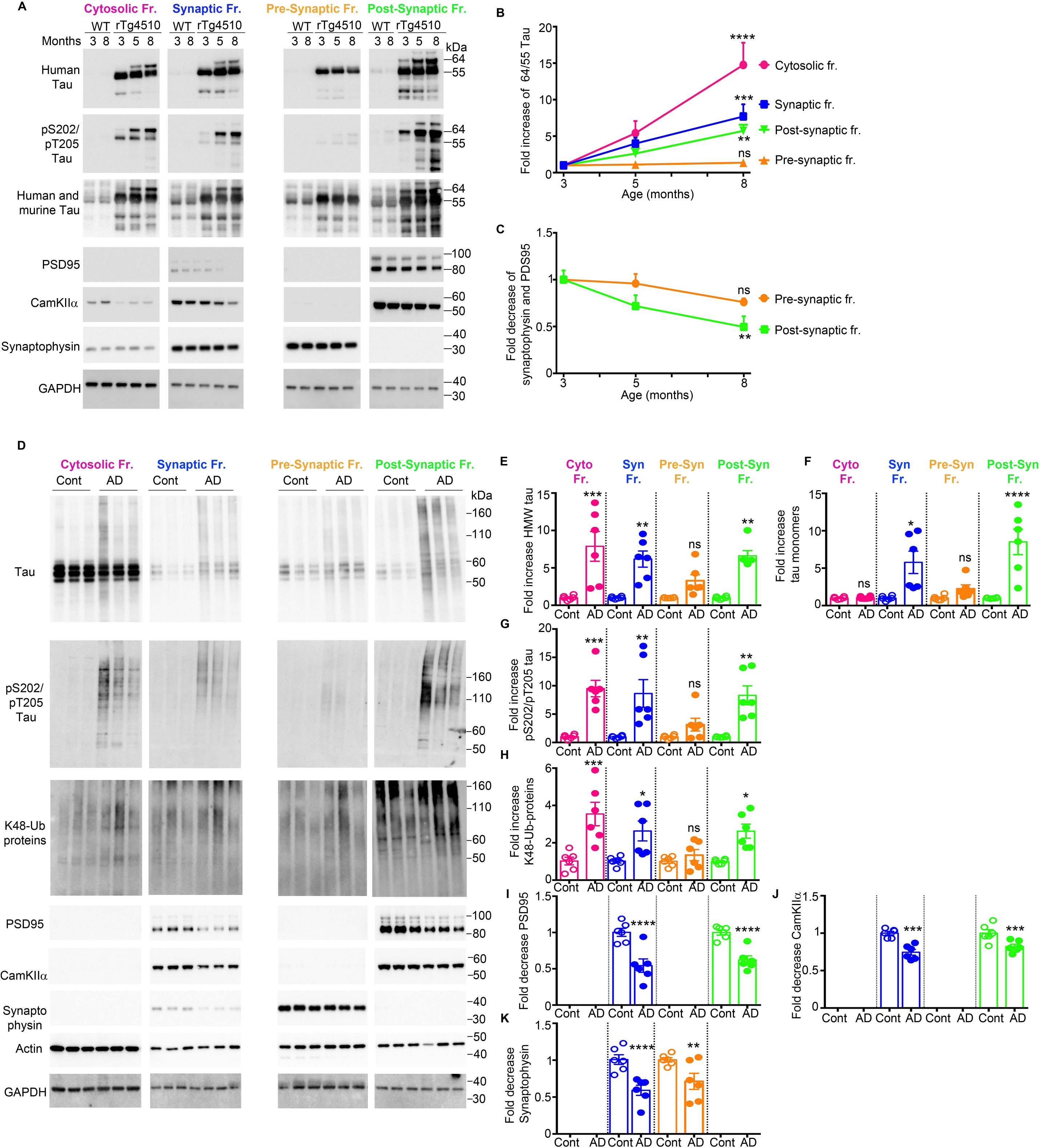
Deposition of pathological tau in the post-synaptic compartments. (**A**) Representative immunoblots of cytosolic, synaptic, pre-synaptic and post-synaptic fractions from WT (3 and 8 months of age) and rTg4510 (3, 5 and 8 months of age) mice for total tau and pS202/pT205 (AT8) tau. Post-synaptic fractions were identified by markers: PSD95 and CamKIIα. Pre-synaptic fractions were identified by synaptophysin. GAPDH was a loading control that was used for normalization. (See Supplementary Fig 1B for uncut immunoblots) (**B**) Quantified densitometry of 64/55-kDa tau ratio, expressed as fold increase relative to 3 months of age. (**C**) Quantified densitometry of pre (synaptophysin) and post (PSD95) -synaptic markers expressed as fold decrease relative to 3 months of age. (**D**) Representative immunoblots of cytosolic, synaptic, pre-synaptic and post-synaptic fractions from human post-mortem brains of AD and age-matched normal brains for total tau, pS202/pT205 tau and K-48 ubiquitinated proteins. Post and pre -synaptic markers were identified by immunoprobing with their respective markers: PSD95 and CamKIIα and synaptophysin. Actin and GAPDH as a loading control. (**E-K**) Densitometric quantification of immunoblots in D expressed as fold change relative to control brains. (See Supplementary Fig. 3 for uncut immunoblots). For **A**, at least three biological experiments were performed. For each fractionation (biological) experiment, three cortices of mice were pooled together, n = 9 cortical brains/age group were used. For **D**, six blocks of BA9 region from AD and age-matched normal post-mortem tissue were used. Error bars, mean ± SEM; n.s., not significant; *P < 0.05, **P < 0.01, ***P < 0.001, ****P < 0.0001. Two-way repeated-measures ANOVA with post hoc Bonferroni correction was used in **Fig.1 B, C**. One-way ANOVA followed by Bonferroni multiple comparison post hoc test was used in **Fig. 1E-K**.

The biochemical state of tau species across fractions was assessed by quantitative immunoblotting, and the ratio of 64-kDa to 55-kDa tau bands (referred to as the 64/55-kDa tau ratio) was used to indicate the tauopathy stage in these mice. The slower migrating, 64-kDa-tau, represents disease-associated, hyper-phosphorylated, and highly aggregable species of tau, whereas the 55-kDa tau is associated with soluble and the physiological form of tau (**Fig. 1A**).

Here we show that cytosolic fraction displayed a 15-fold increase in the ratio between 64/55-kDa tau as tauopathy progressed (from 3-8 months) due to an inverse correlation of 64 and 55-kDa tau levels (**Fig. 1A, B**). In synaptic and post-synaptic fractions, we detected a lower (∼ 6 fold) increase in the 64/55-kDa tau ratio across tauopathy stages due to the parallel accumulation of 55 and 64 -kDa synaptic tau (**Fig. 1 A, B**). Within synapses, pre-synaptic fractions exhibited remarkably different tau composition compared to post-synaptic fractions as levels of 55-kDa and 64-kDa tau progressively increased in post-synaptic fractions from 3 to 8 months old animals, whereas pre-synaptic fractions contained mainly 55-kDa tau with moderate decrease of tau levels from 3 to 8 months in rTg4510 mice (**Fig. 1 A, B**). The latter could be due to retraction of tau from pre-synaptic space and axonal projections to somato-dendritic compartments and/or reduced pre-synaptic terminals.

Next, we assessed AT8 (pS202/pT2015 tau) immunoreactivity across the fractions as this epitope is considered to be a typical marker of early to moderate stage of tau pathology with immunoreactivity confined to soma and dendrite (*77*). Here we show that AT8 positive tau is present in cytosolic and synaptic fractions (**Fig. 1A**). However, within synapses, it is overwhelmingly present in post-synaptic compartments. Moreover, in pre-synaptic compartments, AT8 positive tau is virtually undetectable similar to fractions from WT animals (**Fig. 1A**). This is in accordance with other published studies that have shown that tau accumulates in dendritic spines and that aberrant phosphorylation plays a critical role in tau mislocalization to post-synaptic compartments (*14, 68*). In WT animals, tau displayed equal distribution of synaptic tau in the pre and post -synaptic fractions, and age (3, 8 months) did not affect the sub-synaptic distribution of tau (**Fig. 1A**). Markers for pre-synaptic (synaptophysin) and post-synaptic (PSD95 and CaMKIIα) fractions were used to validate the corresponding fractions, showing a significant reduction of post-synaptic markers in rTg4510 mice, indicative of spine loss (**Fig. 1 A, C**). Synaptophysin exhibited a non-significant reduction in presynaptic fractions, as was reported by other studies in rTg4510 (*70*) (**Fig. 1 A, C**).

To investigate the aggregation state of tau in synapses, we generated tau-enriched insoluble extracts from the total lysate, synaptic, and cytosolic fractions of rTg4510 mice with early, mid and late -stage tauopathy (**Supplementary Fig. 1C**). Our data show that insoluble synaptic tau was present at high levels in two forms, as non-phosphorylated 55-kDa and as phosphorylated (pS202/pT205) 64-kDa tau (**Supplementary Fig. 1 C, D**). Insoluble total extract, similar to insoluble cytosolic extract, contained mainly 64-kDa phosphorylated (pS202/pT205) tau (**Supplementary Fig. 1 C, D**) resulting in 3-fold increase in the ratio of 64/55kDa tau from 3 to 8 months of age, compared to lower (1-fold) increase in 64/55kDa tau ratio in the synaptic fraction (**Supplementary Fig. 1 C, D**). These results suggest that as tauopathy worsens, the synapses are enriched with distinct forms of tau aggregates compared to tau aggregates in other subcellular compartments.

Additionally, we performed a quality control experiment to confirm that the tau observed in post-synaptic fractions was truly accumulated in synapses and was not a contaminant that had co-sedimented with synaptosomes during the sucrose gradient centrifugation. Crude synaptosomes generated from WT mice were mixed with insoluble tau aggregates generated from equal amounts of brain lysate from 3 and 8 months old rTg4510 mice, and together with rTg4510-derived crude synaptosomes were further subjected to discontinuous sucrose gradient centrifugation to isolate purified synaptic fractions (**Supplementary Fig. 2A** (schematic diagram)). Post-synaptic fractions from WT mice mixed with aggregates of 3 and 8 months old rTg4510 mice were either negative or showed negligible amounts of tau, respectively (**Supplementary Fig. 2B**), whereas tau in post-synaptic fractions from rTg450 extracts showed a similar synaptic distribution as in **Fig. 1A**. Input extracts from WT mice mixed with tau aggregates before gradient centrifugation show a similar distribution of tau as extracts from rTg4510 mice (**Supplementary Fig. 2C**).

Furthermore, sedimentation of isolated synapses by continuous glycerol density centrifugation (10-40% glycerol) from rTg4510 and WT mice mixed with insoluble tau aggregate from 8 month old rTg4510 (**Supplementary Fig. 2D**) confirmed that virtually no tau was present in eluants from the corresponding synaptic compartment (fractions 6 and 7) of WT mice mixed with tau aggregates. However, in rTg4510 mice, tau eluted in fractions corresponding to synapses (fractions 6 and 7) (**Supplementary Fig. 2E**). Thus, we confirmed that tau detected in the post-synaptic fractions in **Fig. 1A** represents accumulated synaptic tau and not tau aggregates from other cell compartments.

To examine the accumulation of tau into synapses and the distribution of tau between pre and post -synaptic compartments in AD and age-matched non-demented control brains, we performed a similar fractionation assay using post-mortem brain tissue from cortical Brodmann area 9 (BA9), an area that is progressively affected in AD (*5*) (**Supplementary Table 1**). In patients with AD, tau accumulates into high molecular weight (HMW) oligomers that assemble into filaments and tangles as the disease progresses (*78, 79*). Our data show that cytosolic and synaptic fractions of AD brains exhibit ∼7-10-fold increase in HMW and AT-8 positive tau compared to control cases which display non-phosphorylated monomeric tau enriched in cytosolic fractions and low amount of tau present in synapses (**Fig. 1 D-G**. For uncut blots see **Supplementary Fig. 3**). Moreover, post-synaptic fractions from AD brains accumulate significantly higher levels of monomeric, HMW and AT8 positive tau species compared to pre-synaptic fractions (**Fig. 1 D-G**), consistent with results from rTg450 mice. Previous studies from AD synaptosomes have also reported a trend in the increase in levels of tau in post-synaptic compared to pre-synaptic fraction, albeit using a different approach whereby synaptosome structures were fixed onto a glass slide followed by immunostaining and optical imaging (*11*). As expected, normal brains exhibited a negligible amount of tau in synaptic compartments and a non-significant difference in the distribution of tau between pre and post - synaptic compartments (**Fig. 1 D-F**).

Furthermore, we show that subcellular compartments that exhibited a significant increase in the levels of HMW, AT-8 positive tau species also exhibited a substantial increase in the levels of K48-linked ubiquitin chains that are the most prevalent proteasome-targeting signal (**Fig. 1 D, H**), confirming previous evidence that tau aggregates can cause dysfunction in the UPS-mediated protein clearance (*29, 39, 40*).

Consistent with previous studies (*80*), protein markers of post-synaptic (**Fig. 1 D, I, J**) and pre-synaptic compartments (**Fig. 1 D, K**) were significantly reduced in AD compared to normal brains tissue.

Altogether, our results show that the accumulation of hyper-phosphorylated and aggregated tau into synaptic compartments is a pathological feature of tauopathy disorders. Moreover, we show that missorting and the progressive deposition of pathogenic forms of tau predominantly into post-synaptic compartments occurs in a mouse model of tauopathy and in the brains of AD patients making these synaptic structures particularly vulnerable to tau toxicity. Post-synaptic tau toxicity could be further exacerbated by apparent impaired UPS clearance, which we detected by measuring the levels of K48-linked ubiquitin chains that target proteins to proteasomal degradation.

### Post-synaptic tau species exhibit high seeding activity

An essential feature of tauopathy is that seed competent tau is preferentially transported trans-synaptically across anatomically connected regions (*6, 18, 19, 28, 81*). For trans-synaptic transmission to occur, tau seeds would have to be present in the pre and post -synaptic compartments of neurons and be able to corrupt tau monomers in recipient synapses/cells. Accordingly, we show in Fig.1 that pathological tau can be found in isolated pre and post -synaptic fractions from AD post-mortem brains. Other immunohistological studies have shown tau to co-localize with both pre and post -synaptic markers of isolated synaptosomes from AD patient brains (*11, 29, 82*).

To test whether tau species detected in pre and post -synaptic fractions from mice and human samples have seeding activity, i.e., the ability to recruit and misfold endogenous tau monomers, we used HEK 293 cell line that stably expresses repeat domain of tau with two diseases associated mutations (P301L and V337M) fused with YFP (RD-P301L/V337M-YFP) (*24, 25*). This cell line, (denoted as DS1 clone, a kind gift from Dr. Diamond) lacks tau aggregates, however, upon exposure to exogenous tau seeds, DS1 cells can indefinitely propagate tau aggregation. Previous studies with similar biosensor cells have shown seeding activity of tau from the total (*83*) and synaptosome extracts (*65*) of AD brains, indicating that seeding activity can be a marker of disease progression that manifests before histopathological detection of NFTs. DS1 cells were exposed overnight to an equal amount of tau (3 ng/well) from rTg4510-derived pre and post-synaptic fractions from mice with early, mid and late -stage tauopathy. Pre-synaptic tau exhibited negligible seeding activity across disease stages (**Fig. 2 A, B**), whereas tau from post-synaptic fractions showed significant seeding activity between early and late stages of tauopathy (**Fig. 2A, B**). In contrast, pre and post -synaptic fractions from WT animals showed no seeding activity (data not shown).

**Figure 2.**
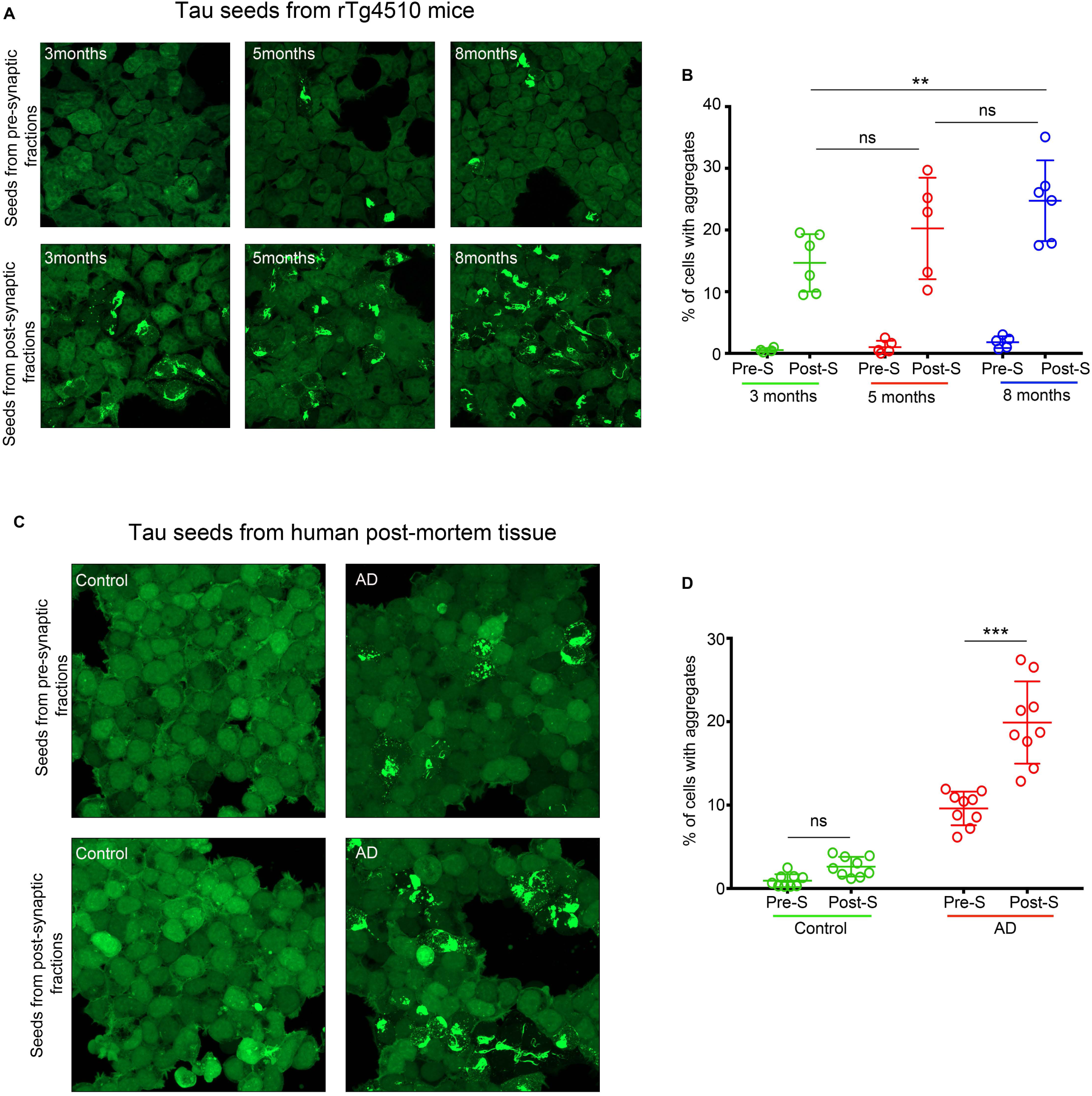
Post-synaptic tau species exhibit high seeding activity. (**A**) Representative images of HEK239-tau RD-P301L/V337M-YFP (DS1) cells exposed to 3ng of tau/well from pre and post -synaptic fractions of rTg4510 mice (3, 5 and 8 months of age). (**B**) Quantification of seeding activity in DS1 cells in A expressed as % cells with aggregates/total number cells (**C**) Representative images of HEK239-tau RD-P301L/V337M-YFP (DS1) cells exposed to 5ng/well of tau from pre and post-synaptic fractions of normal and AD post-mortem brains. (**D**) Quantification of seeding activity in DS1 cells in C expressed as % cell with aggregates over a total number of cells. Scale bars represent 50 μm. Seeds from three different biological extracts were used to test seeding in six (Fig 2A, B) or eight (Fig 2 C, D) independent experiments. *Data* were plotted as mean ± SEM. n.s. not significant; *P < 0.05, **P < 0.01, ***P < 0.001. For statistical analysis, we employed two-way repeated-measures ANOVA with post hoc Bonferroni correction.

Furthermore, when we tested the seeding activity of tau (5ng/well) derived from isolated pre and post - synaptic fractions from normal and AD cases, we observed similar results as with transgenic tau mouse seeds. Pre and post –synaptic tau from normal brains exhibited negligible seeding activity. However, tau from pre and post -synaptic fractions of AD cases displayed significant seeding activity. Specifically, tau from post-synaptic compartments showed significantly more seeding activity compared to tau from pre-synaptic fractions when an equal amount of tau was added to cells (**Fig. 2 C, D**).

These results show that there is a positive correlation between the spatial distribution of tau within synapses and the seeding competency of synaptic tau species to induce aggregation in the biosensor assay. Moreover, seed competent tau was present in post-synaptic compartments before any detectable deficit in post-synaptic markers and before mature tangles were detected in rTg4510 mice.

### PACAP can attenuate seed-induced misfolding of endogenous tau

Upon uptake by neurons, extracellular tau seeds can trigger misfolding and mislocalization of endogenous monomeric tau (*84, 85*). It is proposed that tau seeds can act as a template for the amplification of pathogenic tau, initially within recipient neurons followed by propagation of seeds to neighboring neurons across synaptic connections throughout the brain.

For effective therapies that target the trans-synaptic spread of pathogenic tau, it is vital to target toxic tau in synaptic compartments. One strategy would be to prevent tau aggregates from being released and spread by enhancing tau clearance at the synapses.

We first tested whether stimulation of PAC1R signaling and enhanced proteasome activity by PACAP can lead to reduced misfolding of endogenous tau upon exposure to tau seeds in primary neuronal cultures from the PS19 tau transgenic line that encodes the P301S tau mutation.

After being released from axon terminals, PACAP exerts its physiologic function by binding to, and stimulating PAC1R, a Gs-coupled GPCR, primarily located on post-synaptic membranes of neurons leading to potent activation of cAMP/PKA signaling (*58, 59*). Through these intracellular signaling pathways, PACAP promotes neurogenesis and differentiation during neurodevelopment (*86, 87*), inhibits apoptosis (*88, 89*) and provides neuroprotection (*60, 90–92*) under various toxic stimuli in the developed brain.

PS19 primary neuronal cells were exposed for five days to tau seeds generated from HEK 293 clonal line expressing RD-P301L/V337M-YFP that show tau aggregation and high seeding activity (DS9 clone. A kind gift from Dr. Diamond). The accumulation of conformationally altered and misfolded insoluble tau was assessed after soluble tau was removed by incubating neurons with 0.1% Triton-X100 during the fixation step, followed by MC1 and MAP2 (dendritic marker) mAb immunostaining (*93*). PS19 primary neurons not exposed to seeds (**Fig. 3A**) or treated with PACAP alone (**Fig. 3B**) do not accumulate MC1 positive tau. However, neurons exposed to tau seeds show seeding activity and accumulation of insoluble tau mainly in MAP2 positive dendrites (**Fig. 3C**). One day after seed exposure, neurons were treated for four days with 100 nM PACAP (**Fig. 3D**) or pre-treated with 200nM of a PAC1R antagonist, the N-terminally truncated PACAP(6–38) (*94*), 6 hours before the addition of 100 nM PACAP (**Fig. 3E**). Treatment with PACAP resulted in a significant reduction in the templating of endogenous tau into insoluble MC1 positive tau (**Fig. 3 D, F**), and pretreatment with PACAP antagonist prevented the effect of PACAP treatment alone (**Fig. 3 E, F**). The same experimental paradigm was used when 0.1% Triton-X100 was omitted during the fixation step to show that survival and neurite integrity was not compromised during seed exposure (**Supplementary Fig. 4 A-E**) and during the treatment paradigm (**Supplementary Fig. 4 C-E**). Overall these results suggest that PACAP, via its receptor-PAC1R, mediated intracellular signaling leading to reduced seed-competent tau and the accumulation of insoluble MC1 positive tau, suggesting that targeting tau clearance in dendrite/post-synaptic compartments could be used as a synaptic therapy approach.

**Figure 3.**
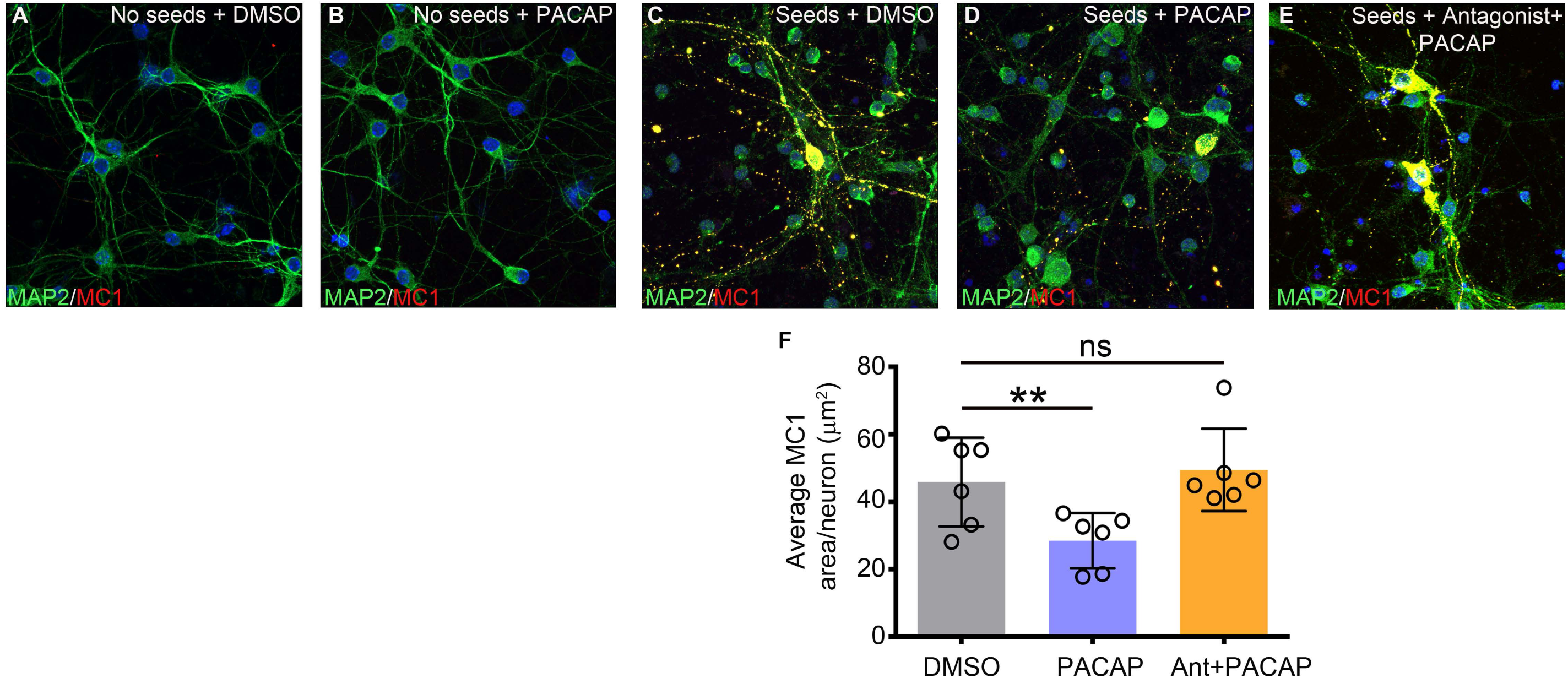
Seed-induced misfolding of endogenous tau can be attenuated by PACAP. Representative images of PS19 primary neuronal cultures treated with (**A**) DMSO or (**B**) PACAP for four days, followed by fluorescent immunostaining. Representative images of PS19 of primary neuronal cultures seeded with 20 µg of DS9 cell lysate (**C-E**) followed by four days treatment with (**C**) DMSO, (**D**) 100 nM PACAP or (**E**) pre-treated with 200nM PAC1R antagonist (PACAP 6-38) for 6 h before adding 100 nM PACAP. The media with PACAP was replenished every 48 hours. Cells were fixed five days after seed exposure. During fixation, 0.1% Triton X-100 was used to remove soluble tau, followed by immunostaining for conformationally altered tau (MC1) and MAP2. (**F**) Quantification of the area of MC1 positive neurons reported as µm^2^ over the numbers of neurons (MAP2 and DAPI positive) per well. Scale bars represent 50 μm. Seeds from three batches of DS9 extracts were used to test seeding in six independent experiments*. Data* were plotted as mean ± SEM. n.s. not significant; *P < 0.05, **P < 0.01. For statistical analysis, we employed one-way ANOVA followed by Bonferroni multiple comparison *post hoc* test.

### PACAP treatment reduces synaptic tau pathology in rTg4510 mice

In situ hybridization studies of adult brains show that PACAP and PAC1R receptors are widely distributed, with the highest amounts of PACAP detected in the hypothalamus (*95, 96*). However, high levels of PACAP and PAC1R are also present in the cerebral cortex, the dentate gyrus, CA1, and CA3 pyramidal neurons of the hippocampus, striatum, and nucleus accumbens (*97–101*). Immunohistochemical labeling of the PAC1R protein shows that, in neurons, the receptor is primarily located in neuronal perikarya and dendrites (*95*). It has been reported that PACAP but not PAC1R levels start to decline before the onset of AD dementia as early as the MCI-AD stage and that this reduction in PACAP levels was shown to be region-specific, targeting vulnerable areas of the AD brain (*102*). Interestingly, a deficit in PACAP was also reported in PS1/APP/tauP301L triple transgenic mice (*103*), and intranasal administration of PACAP in APP transgenic mice showed a reduction of Aβ plaques and increase of sAPPα (*103*). This effect of PACAP on the amyloid pathway could be due to the overall impact of Gs-coupled GPCRs that have been shown to promote the non-amyloidogenic cleavage of APP by α secretase, generating a neuroprotective sAPPα peptide and at the same time reducing Aβ plaque burden (*104, 105*).

Because PACAP, through its receptor, would primarily exert its effect on dendrites and dendritic spines, we used rTg4510 mice known to accumulate tau in post-synaptic compartments early in the disease (**Fig. 1A**), to test whether stimulation of PAC1R receptor by PACAP could promote compartment-restricted activation of PKA/proteasome-mediated clearance of toxic tau species in early-stage (∼4 months of age) rTg4510 mice. Our published studies and others have shown that enhanced proteasome activity through PKA can facilitate the clearance of tau *in vitro* (*106, 107*) and *in vivo* (*40*). In our studies, treatment was achieved by intraventricular administration of PACAP via Alzet osmotic pump, allowing for slow and continuous infusion for 30 days of PACAP (10 pmol/h; with a rate of 0.25 µl/h) and vehicle (0.9% saline and 0.1% BSA). After treatment and behavioral testing, cortical brain regions were harvested and subjected to the sub-cellular fractionation method. Isolated fractions (cytosol, crude synaptosome, synaptosome, pre and post -synaptic fractions) were analyzed by quantitative immunoblotting for tau species.

Our results show that PACAP treatment caused a significant decrease in the levels of total tau in the post-synaptic fraction compared to vehicle-treated mice (**Fig. 4 A, B**. For uncut blots see **Supplementary Fig. 5**). Moreover, the subcellular distribution of phospho-tau species varied between fractions. For instance, pS202/T205 tau, predominantly found in the cytosolic and post-synaptic fractions was significantly reduced in PACAP treated animals, (**Fig. 4 A, C**) and pS396/PS404 tau that had been evenly distributed across all fractions was markedly decreased in the synapses (pre and post -synaptic fractions) (**Fig. 4 A, D**). We also assessed the level of pS214 tau, which is a marker of stimulated cAMP/PKA signaling and an indication that PKA signaling was active during PACAP treatment. In the cytosolic fraction, where most metabolic chemical reactions occur, we detected a significant increase in pS214 tau levels in PACAP treated compared to vehicle-treated littermates (**Fig. 4 A, E**). Within synapses, pS214 tau was predominately found in the pre-synaptic compartment and was significantly reduced in PACAP treated mice (**Fig. 4 A, E**). Interestingly, the migration pattern of 55 and 64 kDa tau species appeared to differ between pre and post - synaptic compartments, with pre-synaptic fractions containing 55-kDa tau and post-synaptic fractions containing predominately the 64-kDa tau forms, which is recognized to be a pathological and aggregated form of tau (**Fig. 4A**).

**Figure 4.**
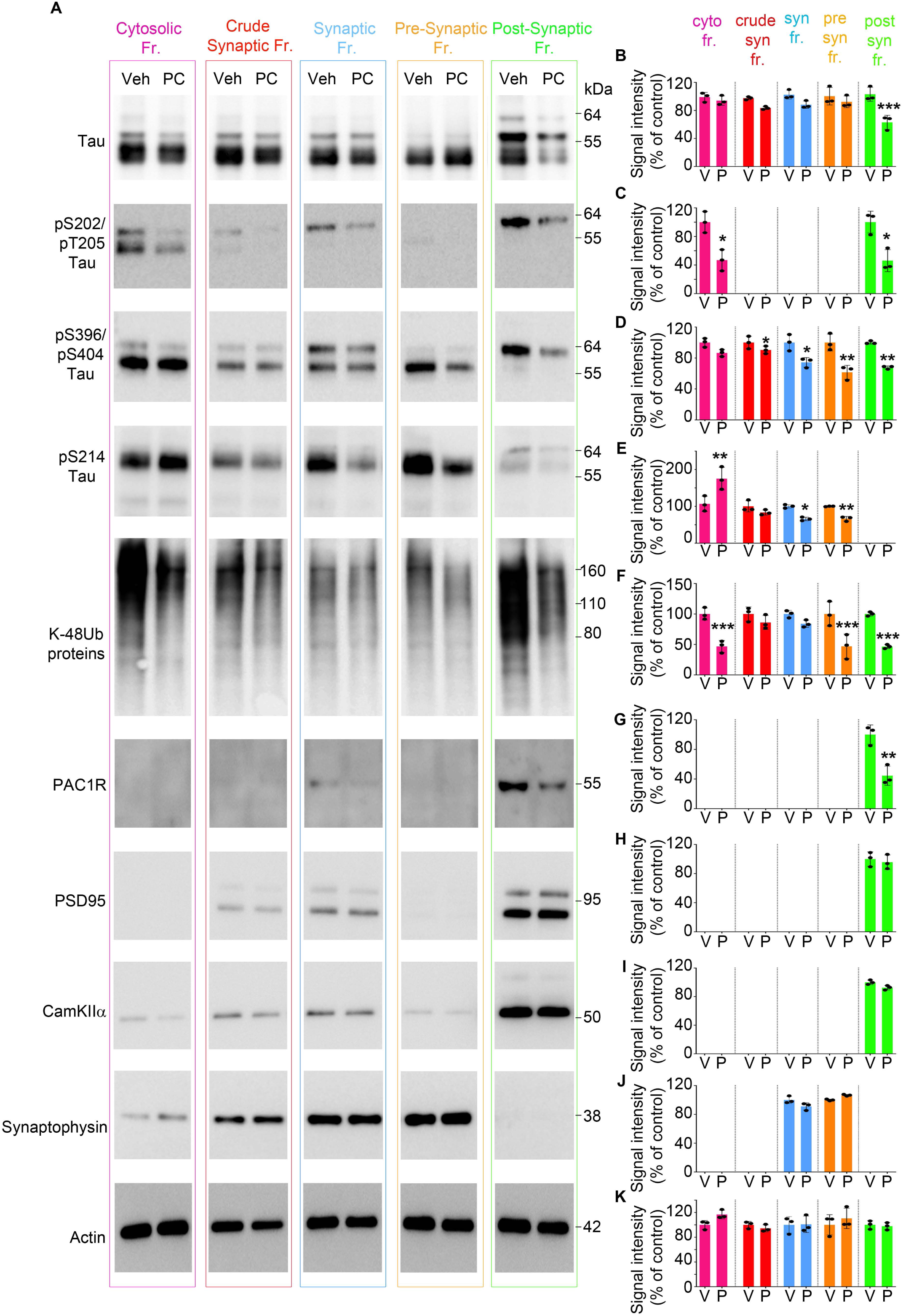
PACAP treatment reduces synaptic tau pathology in rTg4510 mice. (**A**) Representative immunoblots of cytosolic, crude synaptic, synaptic, pre-synaptic and post-synaptic fractions from rTg4510 mice treated with vehicle or PACAP. Blots were immunoprobed for total tau and phospho tau species (pS202/pT205, pS396/pT205, and pS214 tau epitopes) and PACAP receptor - PAC1R. Post-synaptic fractions were identified by markers: PSD95 and CamKIIα. Pre-synaptic fractions were identified by synaptophysin. Actin was a loading control that was used for normalization. (**B-K**) Quantified densitometry of A, expressed as fold change relative to vehicle-treated. At least three biological experiments were performed. For each fractionation experiment, 5 hemi cortices were pooled together for fractionation experiment, n = 15 hemi cortical brains/treatment. Error bars, mean ± SEM.; *P < 0.05, **P < 0.01, ***P < 0.001. For statistical analysis, we employed one-way ANOVA followed by Bonferroni multiple comparison post hoc test.

Next, we tested the amount of K48-linked poly-ubiquitinated proteins across fractions to determine proteasome activity indirectly. Our data show that proteasome-specific ubiquitinated proteins were significantly lower in cytosolic, pre and post-synaptic fractions (**Fig. 4 A, F**), suggesting enhanced proteasome activity resulted in reduced ubiquitinated protein levels.

Moreover, we tested the distribution and the levels of PAC1R receptors upon PACAP treatment. We show that the PAC1R receptor is present predominantly in the post-synaptic fractions, confirming what other studies have suggested that the receptor is present on the membrane of dendrites (**Fig. 4 A, G**). Furthermore, PACAP treatment caused a significant decrease in the levels of PAC1R in the post-synaptic compartments (**Fig. 4 A, G**), showing target engagement due to the effect of agonist-induced activation of the receptor leading to internalization for fine-tuning of the PAC1R signaling (*108–110*).

Extracts from fractionation assays were tested for markers of post-synaptic (PSD95 and CamkII) (**Fig. 4 A, H, I**) and pre-synaptic (Synaptophysin) (**Fig. 4 A, J**) compartments, which were detected in the corresponding fractions. Their levels were not changed upon PACAP treatment.

Overall the results show that *in vivo* stimulation of a GPCR (PAC1R receptor) by its ligand can propagate intracellular signaling that can lead to reduced total tau in the post-synaptic compartment and reduced phospho-tau in synapses and overall reduced UPS-specific ubiquitinated proteins in rTg4510 mice.

### PACAP-via PKA increases phosphorylation and the activity of synaptic 26S proteasome

In previous studies, we have shown that increasing cAMP/PKA signaling by PDE inhibitors resulted in PKA-mediated phosphorylation and enhanced proteolytic activity of 26S proteasomes leading to clearance of mutant tau and overall attenuation of tauopathy in mice (*40, 48*).

Because PACAP-induced signaling is expected to be restricted to where the PAC1R receptor is present, we hypothesized that PACAP treatment would enhance proteasome function predominantly in post-synaptic compartments. Published studies on the subcellular distribution of 26S brain proteasomes have shown that UPS components are abundant in synapses and there are no detectable differences in proteasome subunit composition between cytosolic and synaptic proteasomes (*111*). To test whether PACAP preferentially enhanced synaptic proteasome activity, we isolated synaptic and cytosolic 26S proteasomes by the affinity purification method, which yields purified and functionally intact 26S proteasomes (*40, 112*).

First, we performed a kinetic assay, whereby degradation of a fluorogenic substrate by purified 26S proteasomes was monitored over 60 min (**Fig. 5A**). As shown, PACAP treated 26S proteasomes displayed a higher degradation rate of the fluorogenic substrate compared to vehicle-treated 26S proteasomes (**Fig. 5B**). Specifically, PACAP-treated synaptic 26S proteasomes showed a significantly higher slope of reaction compared to PACAP treated-cytosolic 26S proteasome (**Fig. 5B**). Moreover, purified 26S proteasomes were resolved by native PAGE assay to assess the in-gel proteolytic activity of doubly and singly -capped 26S particles (**Fig. 5C**). The singly-capped 26S proteasomes from PACAP treatment displayed a significant increase in activity (**Fig. 5D, E**), with synaptic 26S singly-capped proteasomes displaying the highest (∼2.5) fold increase in activity (**Fig. 5D, E**).

**Figure 5.**
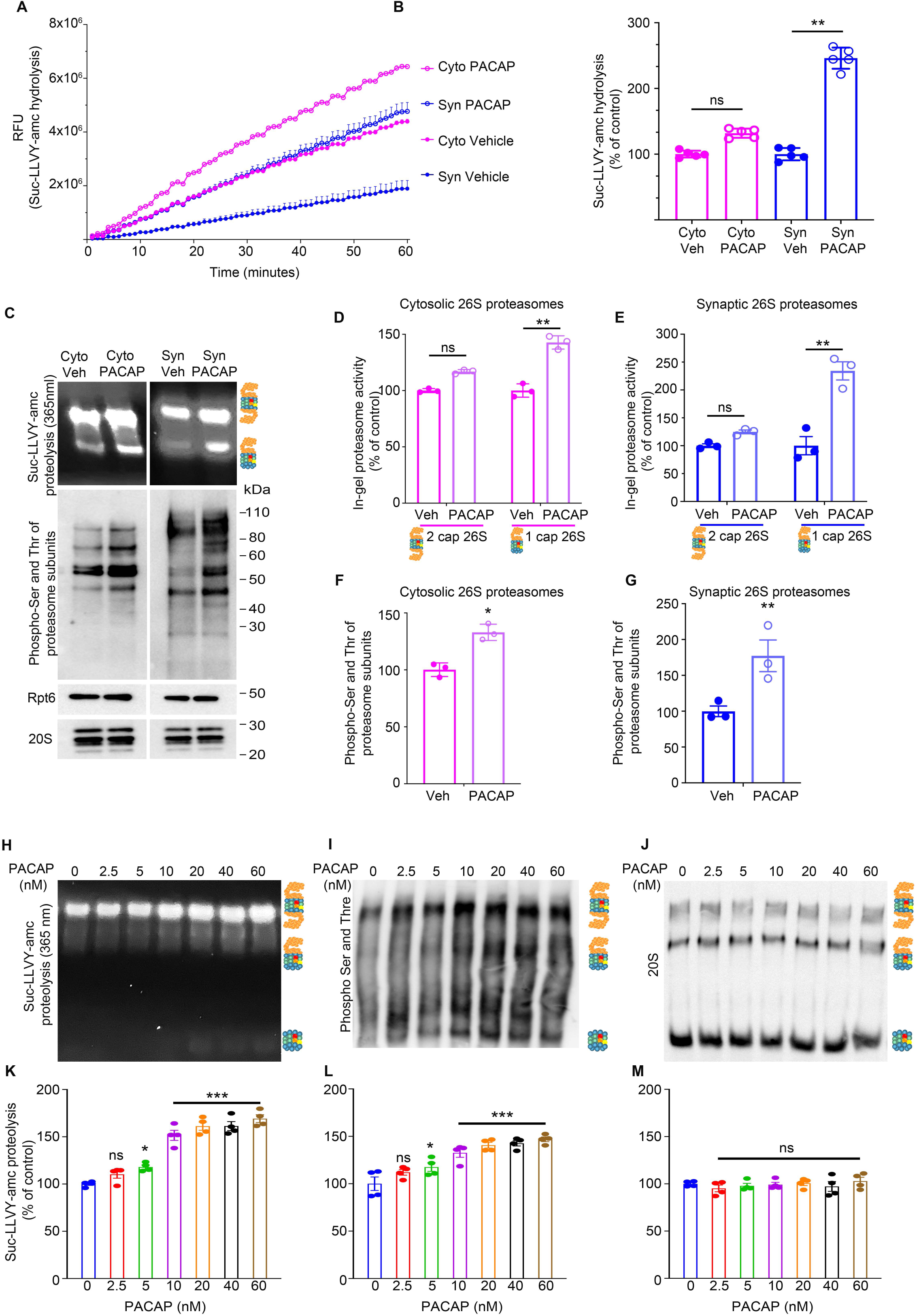
PACAP-via PKA, increases phosphorylation and the function of 26S proteasomes. Purified 26S proteasomes from cytosolic and synaptic fractions of vehicle and PACAP treated rTg4510 mice tested for: (**A**) The peptidase activity, 10 nM of 26S proteasomes were monitored over 60 min using the fluorogenic substrate Suc-LLVY-amc (40 μM). (**B**) Rate of substrate hydrolysis by 26S proteasomes from A. (**C**) 26S purified proteasomes tested for in-gel proteasome activity following native PAGE (top), and immunoblot analysis of PKA-specific phosphorylated serine and threonine (middle), and for proteasome subunits Rpt6 and 20S (bottom). (**D** and **E**) Densitometric quantification of in-gel activity normalized to 26S proteasome levels. (**F** and **G**) Densitometric quantification of PKA-specific phosphorylation of serine and threonine. HEK293 PAC1R-EGFP-expressing cells treated with increasing concentration of PACAP were resolved on a native PAGE for (**H**) In-gel 26S proteasome activity, followed by immunoblotting for (**I**) PKA-specific phosphorylation of serine and threonine and for (**J**) 20S core proteasomes. (**K, L,** and **M**) Densitometric quantification of H, I, and J. Purified 26S proteasomes from *in vivo* study were pooled from n = 6 cortical hemi brains, and at least three independent purification experiments were performed. Data from HEK293 PAC1R-EGFP-expressing cells are representative of at least three independent experiments. Statistical analyses employed one-way ANOVA followed by Bonferroni multiple comparison post hoc test (B, K, L and M), two-way repeated measures of ANOVA with Bonferroni multiple comparison post hoc test (D and E), and two-tailed Student’s t-test between groups (F and G). Error bars, mean ± SEM.; n.s., not significant; *P < 0.05, **P < 0.01, ***P < 0.001.

To assess whether stimulated PKA enhanced proteasome activity via serine and threonine phosphorylation of proteasome subunits we analyzed purified 26S proteasome from fractions by quantitative immunoblotting with an antibody to PKA-dependent phosphorylation (**Fig. 5C**). A significant increase in the phosphorylation of serine and threonine epitopes of several subunits was evident in cytosolic and synaptic fractions of the PACAP-treated group, with a more prominent increase in phosphorylation of PACAP treated synaptic proteasomes (**Fig. 5F, G**). The levels of 26S proteasomes represented by Rpt6 and 20S subunits remained unchanged (**Fig. 5C**).

PACAP-mediated enhanced proteasome activity was confirmed in PAC1R-expressing HEK293 cells (A kind gift from Dr. Victor May). Treating these cells with increasing concentrations of PACAP for 16 hours led to a concentration-dependent increase in activity (**Fig. 5H, K**) and phosphorylation levels (**Fig. 5I, L**), of 26S proteasomes without a change in 26S proteasome levels (**Fig. 5J**).

Overall our data demonstrate that stimulation of PAC1R, present predominantly on the post-synaptic compartment, resulted in a pronounced PKA-dependent phosphorylation and activity of synaptic 26S proteasomes.

### PACAP treatment attenuates tauopathy

Next, we assessed whether PACAP treatment could reduce tau phosphorylation and tau aggregation throughout the cortex. A previous study has shown that intranasal administration of a PACAP analog to spinobulbar muscular atrophy (SBMA) mice, characterized by phosphorylation and polyglutamine expansion of the androgen receptor (AR), reduced phosphorylation and aggregation of the mutant protein (*113*). For our studies, cortices of rTg4510 mice treated with PACAP or vehicle were used to prepare total and insoluble extracts, and the levels of total and phosphorylated tau species were assessed. PACAP treatment significantly reduced amounts of total and phosphorylated tau in both the total extract (**Fig. 6A**) and the insoluble fraction (**Fig. 6B**. For uncut blots see **Supplementary Fig. 6**). Consistent with the data from cytosolic fraction, PACAP treatment increased the levels of pS214 tau in the total extract confirming that PKA activity was enhanced during *in vivo* continuous infusion of PACAP (**Fig. 6A**). Immunohistochemical analyses also show that PACAP treatment reduced levels of pS396/pS404 tau significantly (**Fig. 6C**).

**Figure 6.**
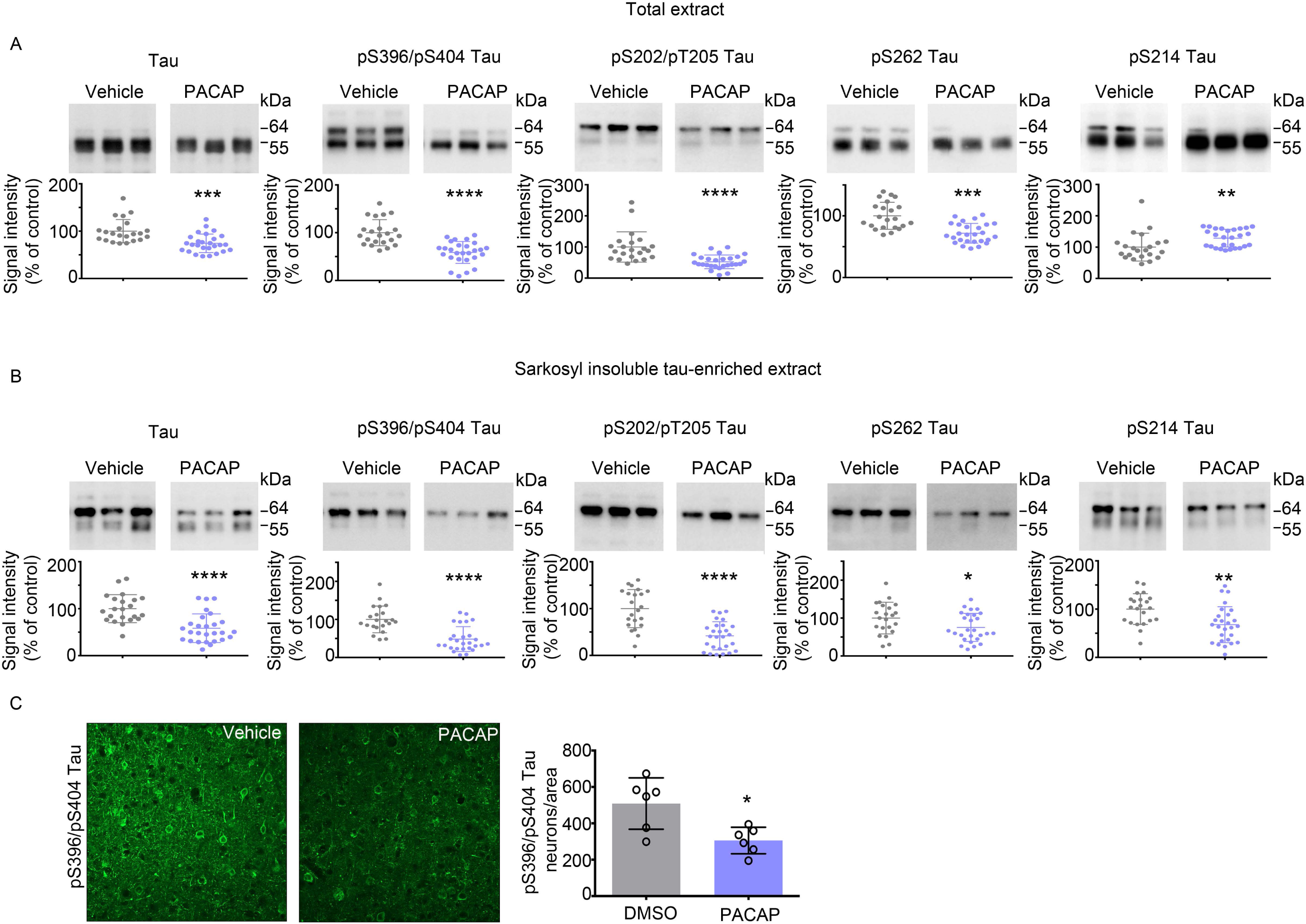
PACAP treatment attenuates tauopathy. Representative immunoblots and corresponding densitometric quantification of (**A**) total and (**B**) insoluble extracts for total tau, pS396 and pS404, pS202 and pT205, pS262, and pS214 tau epitopes from cortical tissue of rTg4510 mice treated with vehicle or PACAP. (**C**) Immunofluorescence labeling and quantification of fluorescence intensity for pS396 and pS404 tau epitope. Scale bars represent 50 μm. Scatter plots represent quantification of immunoreactivity normalized to GAPDH. Statistical analyses of vehicle (n = 22) and PACAP (n = 27) -treated mice were performed in two sets. For quantification of immunofluorescence signal in C, slices from 6 mice per treatment group were analyzed. Error bars, mean ± SEM. *P < 0.05, **P<0.01, ***P < 0.001, ****P<.0.0001. Statistical analysis employed unpaired two-tailed Student’s t-test between two groups.

These results suggest that designing therapies that target clearance of mislocalized post-synaptic tau can attenuate hyper phosphorylation and aggregation of tau throughout the brain.

### PACAP improves cognitive performance in early-stage tauopathy

To assess the effect of PACAP on tauopathy-associated cognitive impairment, we examined WT and rTg4510 mice for hippocampal-dependent spatial learning memory and declarative (episodic) memory, by Morris water maze test (MWM) and novel object recognition test (NOR), respectively. In the MWM test, we observed a difference between PACAP and vehicle-treated rTg4510 mice (treatment effect F [1, 45] = 18.3, p < 0.001) and between WT and rTg4510 mice (genotype effect F [3, 62] = 20.6, p < 0.0001) (**Fig. 7A**). PACAP treated animals showed a significant reduction in latency to reach the hidden platform starting at day 1 of the experiment (t=3.08, df=310, p < 0.05). This effect was sustained at day 2 (t=2.91, df=310, p < 0.05), day 4 (t=3.10, df=310, p < 0.05) and day 5 (t=5.25, df=310, p < 0.001) of the experiment. The effect could not be attributed to a general improvement in cognition or performance, as PACAP treatment did not affect the performance of WT mice (day 5 [t=0.05, df=310, ns]). Summary of behavioral performance is visualized by trace analyses of the swim path in a heat map, showing improvement in latency of PACAP treated rTg4510 mice (**Fig.7 B**).

**Figure 7.**
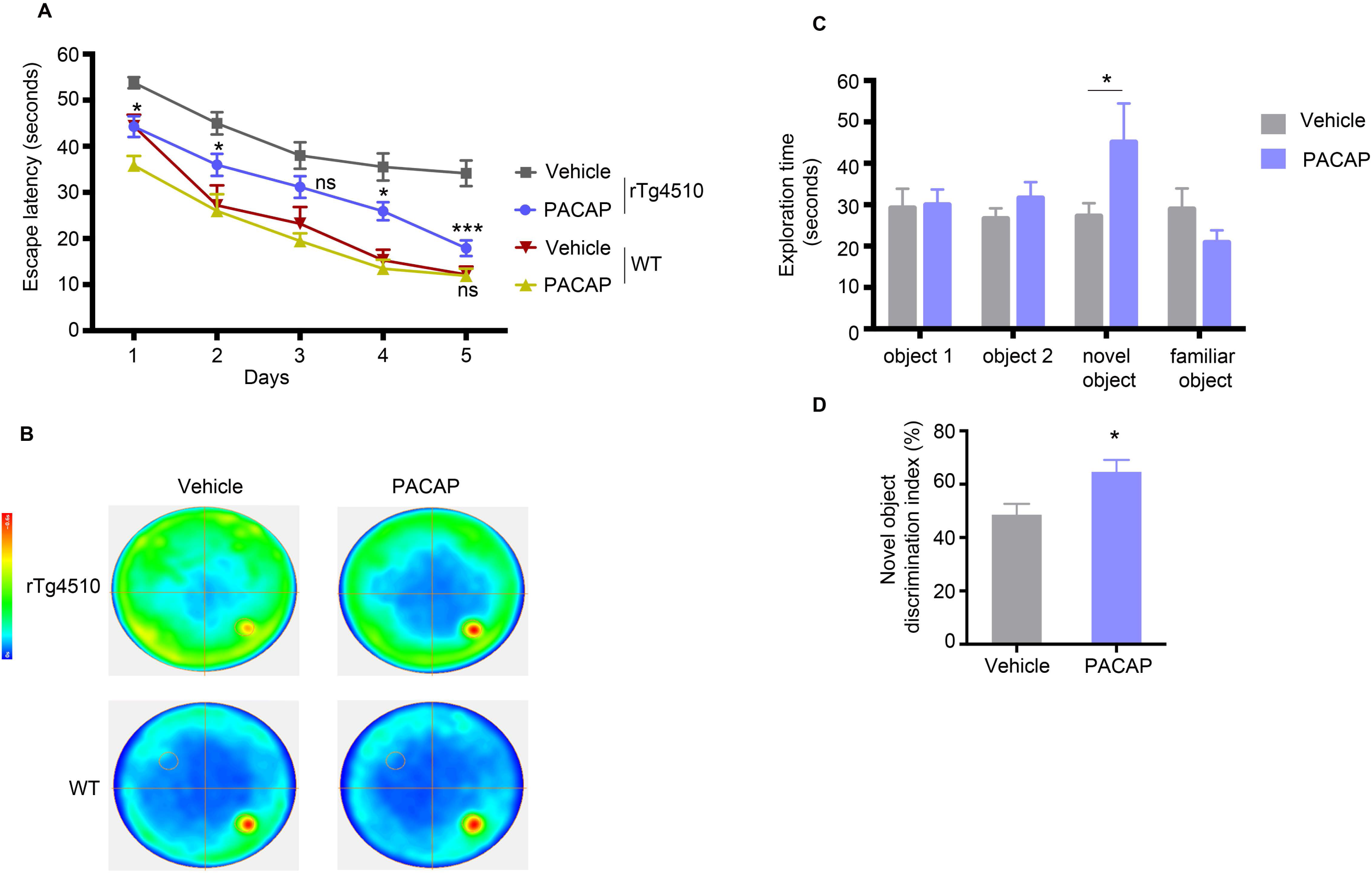
PACAP improves cognitive performance in early-stage tauopathy mice. Morris water maze (MWM) task was used to assess spatial reference memory of rTg4510 and WT mice treated with vehicle or PACAP; n = 22 (vehicle) or 27 (PACAP) rTg4510 mice group, and n = 10 (vehicle) or n =9 (PACAP) in WT mice group. (**A**) Escape latencies were averages of six trials for each day (60 s per trial) separated into two sessions a day (three trials per session). (**B**) Heatmaps depict the summary recording of the MWM task. Increasing color intensity (arbitrary scale) represents increased time spent i.e strong preference for the location where the platform was placed (small circle in the bottom right quadrant). (**C**) Novel object recognition (NOR) task showing exploration, during the object acquisition phase (object 1 and 2) and during object recognition trial (novel and familiar object) for the vehicle (n=22) and PACAP (n=27) treated animals. (**D**) The discrimination index for the novel object was calculated as the difference in time exploring the novel and familiar object, expressed as the ratio of the total time spent exploring both objects (i.e., [Time Novel − Time Familiar/Time Novel + Time Familiar]× 100). Morris water maze and novel object recognition tests utilized two-way repeated-measures ANOVA with post hoc Bonferroni correction. The discrimination index from novel object recognition data was analyzed using unpaired two-tailed Student’s *t*-test between groups with unequal variance. Data are reported as means ± SEM.; n.s., not significant, *P < 0.05, ***P < 0.001.

We next evaluated the exploratory behavior of mice by the NOR paradigm. During the object acquisition phase, mice spent equal time exploring object 1 (t=0.109, df=189) and 2 (t=0.71, df=189) (**Fig. 7C**). During the object recognition trial, PACAP treated mice showed a preference for a novel object, spending significantly more time exploring the novel object (t=2.543, df=189, p<0.05) (**Fig. 7C**). The discrimination index was also increased significantly (t=2.656, df=46, p<0.0108) in PACAP treated littermates (**Fig. 7D**). Vehicle treated mice failed to discriminate between novel and familiar objects, and they spent equal percent time exploring both objects.

## Discussion

Normally tau is a cytosolic protein that is not found in large amounts in synapses (*29*). However, in the early stages of AD, tau is missorted to the dendritic compartment where it is thought to alter synapse stability, contribute to synaptic dysfunction and to loss of dendritic spines (*14, 72, 114–117*). Using a sub-cellular fractionation assay, we examined the biochemical properties of synaptic tau in the pre and post - synaptic compartments across tauopathy stages in the rTg4510 mice and in human control and AD post-mortem brains. Our studies in rTg4510 mice show that highly aggregable (∼ 64kDa) and AT-8 positive tau species were found to accumulate progressively across stages of tauopathy in the post-synaptic compartments, which correlated with a significant decrease in post-synaptic markers. Interestingly, pre-synaptic compartments were moderately enriched with non-phosphorylated tau that remained the same across tauopathy stages. Consistent with the mouse data, HMW, AT-8 positive synaptic tau in the human AD brain accumulated mainly in post-synaptic compartments. Furthermore, synapses from normal brains exhibited low levels of tau with no apparent difference in distribution between pre and post -synaptic compartments. During disease progression, several mechanisms working simultaneously could account for the overwhelming deposition of tau in post-synaptic compartments. For instance: 1) Mislocalization of phospho-tau to dendrites that occurs early in disease pathogenesis. 2) Release of tau from intact or degenerating pre-synaptic terminals into the extracellular space that can be readily taken up by post-synaptic compartments. 3) Misfolded phospho-tau in post-synaptic compartments could impair local proteolysis resulting in failure to remove aggregation-prone tau and proteotoxicity. Our finding that post-synaptic compartments from the AD brain contain high levels of ubiquitinated proteins compared to pre-synaptic compartments supports the third idea. Other published evidence suggests that phospho-tau oligomers present in synaptosomes interact directly with synaptic 26S proteasomes, potentially impeding the proteolytic activity of the local 26S proteasomes causing accumulation of ubiquitinated proteins (*29*). A recent study in microfluidic devices suggests that robust proteolysis of tau in synapses contributes to the polarized distribution of tau mainly in axons and that downregulation of UPS or autophagy in synapses leads to missorting of tau to dendrites followed by loss of dendritic spines (*15*).

The toxic tau species that accumulate in post-synaptic compartments early in the disease can act as seeds for aggregation and spread of tau pathology in other subcellular compartments within a neuron and across synapses to neighboring neurons. Recent studies have demonstrated that seed competent tau that can cause aggregation of naïve monomeric tau is present in the total lysate (*83*) and synaptosome (*65*) extracts of human AD brains before the onset of tau pathology detected by IHC. Herein our results extend these findings by demonstrating that isolated post-synaptic compartments contain tau that exhibits significantly higher seeding activity than tau in the pre-synaptic compartment from transgenic mice and human AD brains. This apparent difference in pathogenicity of tau seeds could be due to the differential distribution of tau species across synapses, i.e., HMW phospho-tau that is overwhelmingly present in post-synaptic compartments. Furthermore, the seeding activity of post-synaptic tau is evident even before overt somatic aggregation of tau in rTg4510.

To develop effective therapeutic strategies that can target accumulation of tau in synaptic compartments in early stages of the disease before the accumulation of neurofibrillary tangles, we employed a strategy of GPCR-mediated stimulation of downstream cAMP/PKA signaling and subsequent enhancement of proteasome-mediated clearance of tau in post-synaptic compartments (**Fig. 8**).

**Figure 8.**
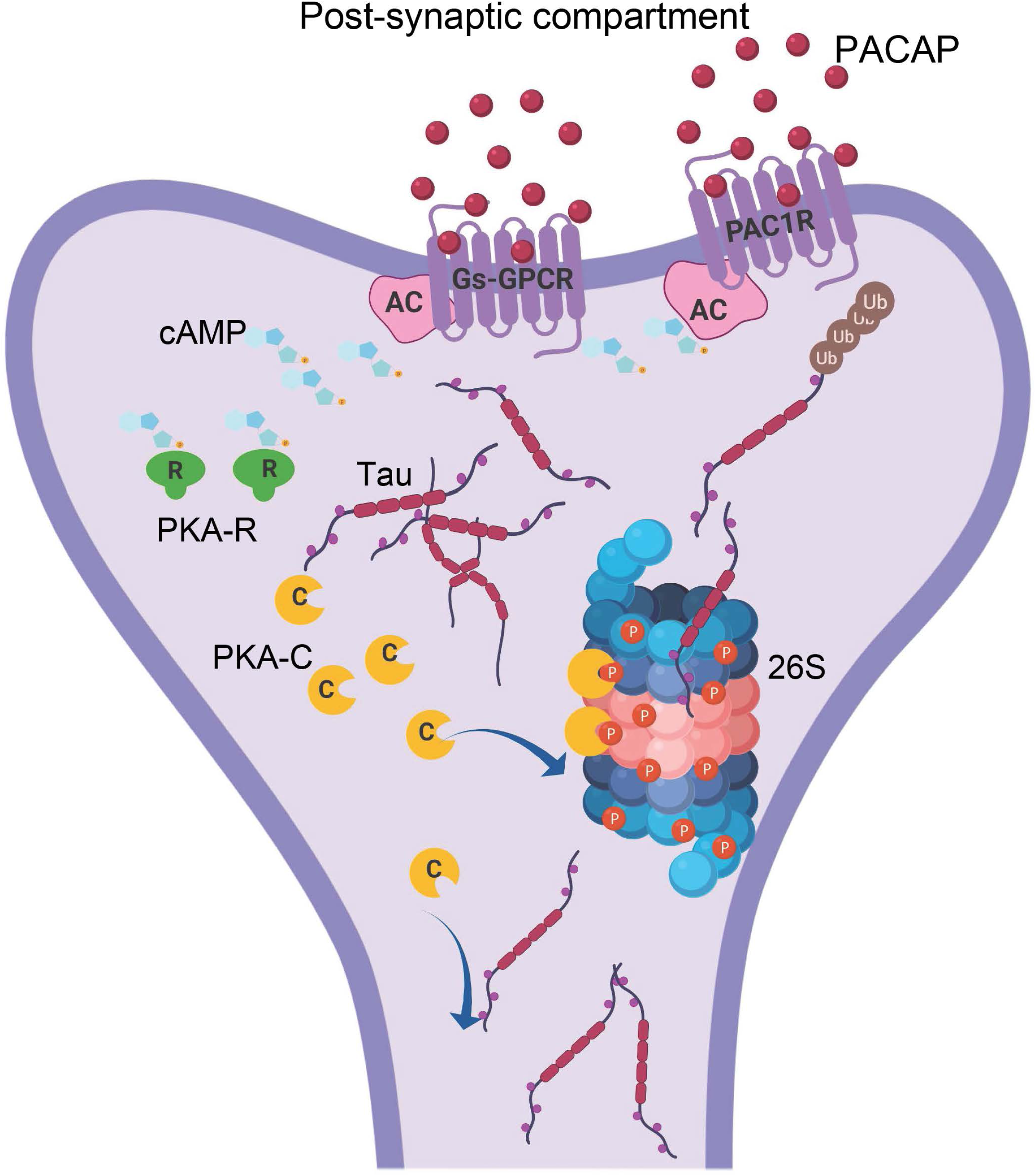
Tau clearance in the post-synaptic compartment. A schematic illustration of how PACAP stimulation of the PAC1 receptor, a Gs GPCR, present on the membrane of the post-synaptic compartment, can lead to activation of AC/cAMP/PKA signaling. Liberated activate PKA can phosphorylate 26S proteasome subunits, conferring super activity on the proteasomes leading to enhanced clearance of missorted tau in the post-synaptic compartment.

The GPCRs mediate critical physiological functions and are considered one of the most successful drug targets, as ∼34 % of all drugs approved by the FDA target 108 GPCRs for a broad spectrum of diseases (*118*). Activation of Gs-coupled GPCRs that transduce the classical AC/cAMP/PKA signaling has been suggested as a therapeutic approach for AD primarily due to its anti-amyloidogenic effect involving alpha-secretase dependent processing of APP and production of sAPPα, which is proven to be neuroprotective (*104, 119*). However, stimulation of Gs-coupled GPCR in tau-related pathology is less explored. Since GPCRs mediate slow synaptic transmission, then specific Gs-coupled GPCRs that are present in post-synaptic and pre-synaptic membrane terminals (*55*) can stimulate cAMP/PKA signaling and presumably enhance UPS-mediated protein clearance in synapses. In our study, we investigated the effect of *in vivo* stimulation of the PAC1R receptor, a Gs-coupled GPCR, present on the membrane of dendrites of neurons in a mouse model that accumulates tau in the post-synaptic compartment in the early stage of the disease. Our data show that PACAP, through PAC1R, mediates therapeutic effect by enhancing PKA-dependent phosphorylation and activation of proteasomes and clearance of tau restricted to post-synaptic compartments (**Fig. 8**). Chronic treatment with PACAP resulted in a significant reduction of phosphorylated tau species in synapses and throughout the brain. We and others have shown previously that enhanced proteasome activity through PKA is dependent on phosphorylation of proteasome subunits (*40, 49*). Here we show that proteasomes purified from synaptic and cytosolic fraction of PACAP treated animals show an increase in proteolytic activity and an increase in phosphorylation pattern of proteasome subunits compared to purified proteasome vehicle-treated animals. Within PACAP treated proteasomes, synaptic proteasomes display more pronounced activity and phosphorylation compared to cytosolic proteasomes. In line with our work, a previous study has shown in a mouse model of HD (lineR6/2) that *in vivo* chronic stimulation of the A2A receptor resulted in a Gs-coupled response, improved proteasome function by PKA-mediated phosphorylation and reduced HTT aggregates in striatal synapses (*120*). A recent study in tau transgenic mice expressing the FTD mutation DK280 demonstrated that administration of rolofylline, an antagonist of the Gi-coupled adenosine A1 receptor located on the membrane of synaptic terminals, which can lead to elevated cAMP signaling, restored neuronal activity, and prevented pre-synaptic impairment and dendritic spine loss. In this study, the status of synaptic proteasome function and synaptic tau concentrations were not assessed, but the same improvements in synapses were seen when tau aggregation was reduced (*121*). Furthermore, restoring dendritic spines loss by elevating cAMP levels by rolipram, a PDE4 inhibitor, was demonstrated to be dependent on activation of the UPS clearance pathway (*122*). Additionally, our results from behavioral analyses that evaluated hippocampal-dependent spatial learning memory and declarative (episodic) memory showed that PACAP treatment leads to significant improvement of cognitive performance.

To address the limitation of this study, it is essential to note that although PACAP is known to have neurogenic, anti-apoptotic and neuroprotective properties, its metabolic instability remains a major issue and represents a limitation in clinical applications. Previously it was reported that PACAP rapidly undergoes enzymatic degradation by endopeptidases after its administration into the systemic circulation (*123, 124*). With the reported half-life of 5-10 min in the blood, PACAP would reach its site of action in the CNS at a diminished concentration (*124*). This disadvantageous property of PACAP reinforces the necessity to design PACAP derivatives or small molecules with chemical modifications that increase their enzymatic stability while maintaining selectivity and potency for the PAC1R receptor. While the clinical utility of PACAP is not supported, this study presents a novel therapeutic approach aimed at reducing post-synaptic tau and attenuating the spread of pathological tau by pharmacological modulation of GPCRs present on the membrane of post-synaptic compartments. Depending on the distribution of GPCR targets, this strategy could be designed to prevent the accumulation of proteins with regional, cellular, and sub-cellular selectivity in the brain.

In conclusion, we show by subcellular fractionation method the biochemical properties of tau in pre and post -synaptic compartments separately that led us to identify the difference in distribution and seeding activity of tau in synaptic compartments in AD and a mouse model of tauopathy. Post-synaptic compartments were shown to contain high levels of pathological tau, and potentially making these structures vulnerable to tau toxicity. To prevent tau toxicity in the post-synaptic compartment, we designed a therapeutic strategy of space-restricted receptor-mediated clearance of synaptic tau via the proteasome pathway, leading to overall reduced tau pathology and improved cognitive performance in the tauopathy mouse model.

## Acknowledgment

We thank Dr. Marc Diamond for providing us the HEK293 RD-YFP (DS1 and DS9 clone) cell lines and Dr. Victor May for providing the HEK 293 PAC1R-EGFP cell line. We thank Dr. Peter Davies for the gift of tau antibodies. We thank Dr. Benjamin Lassus for helping us with ImageJ analysis for the quantification of the MC1 signal in Fig. 3. We thank Dr. Tal Nuriel for helpful discussions throughout the study.

## Author contributions

N.M. conceived the study. N.M. and K.E.D designed the study. N.M. wrote the manuscript. K.E.D. extensively edited the manuscript. N.M. performed most of the experiments and analyzed the data. A.W.S. performed intracerebroventricular administration of alzet pumps, behavioral and primary cell cultures experiments. A.M.R. performed in-gel assays for proteasome activity in PAC1R-expressing HEK293 cells. S.S. provided the PAC1 antibody. S.L.F. and I.S.M. provided human tissue.

## Data and materials availability

All data associated with this study are present in the paper or the supplementary materials.

## MATERIALS AND METHODS

### Antibodies

Monoclonal antibodies to total human tau (CP27; 1:5000), pS396 and pS404 (PHF1; 1:2500), and a conformational monoclonal antibody specific for PHF-tau (MC1; 1:500) were generous gifts from Dr. Peter Davies. Rabbit anti-human tau was from Dako (1:8000, A0024). Other tau antibodies: polyclonal rabbit pS214 tau was from Life Technologies (1:3000, #44-742G); mouse monoclonal pS202 and pT205 Tau (AT8, 1:2500, #MN, 1020) and rabbit polyclonal pS262 (1:2000, #44-750G) were from Thermo Fisher. Monoclonal anti-rabbit K-48 linked ubiquitin (1:1000, D905) from Cell Signaling. Mouse monoclonal anti-PSD-95 (1:5000, 6G6-1C9, #ab2723); rabbit monoclonal synaptophysin (YE269, ab32127, 1:5000) were from Abcam. CamKIIα (1:2000, A-1, sc13141) was from Santa Cruz Biotechnology. Dendritic marker, anti-rabbit MAP2 (1:500, ab32454), was from Abcam. Rabbit polyclonal PAC1R antibody (1:1000) was provided by Dr. Shioda. Rabbit monoclonal phospho-(Ser and Thr) PKA substrates (1:1000, #100G7E) from Cell signaling. Proteasome antibodies: Mouse monoclonal anti-Rpt6/S8 (#PW9265) and anti-proteasome 20S (α1–α7) (clone MCP231, #BML-PW8195) were from Enzo. Loading control markers anti-GAPDH (1:5000, clone GADH-71.1, #G8795) and Actin (1:5000 AC-74, #A2228) were from Sigma. Secondary antibodies were from Jackson Immunoresearch, anti-mouse (115-035-003) and anti-rabbit (111-035-003).

### Animals

#### Mouse models

Protocols and procedures were approved by the Committee on the Ethics of Animal Experiments of Columbia University and were in full compliance with the US National Institutes of Health Institutional Animal Care. Double-transgenic rTg4510 mice express human tau with four microtubule-binding domain repeats (4R0N) and the P301L mutation (line FVB-Tg(tetO-*MAPT*\*P301L human)#Kha/JlwsJ) under the control of mouse calcium-calmodulin kinase II–driven tetracycline-controlled transcriptional activator (tTa) (line Tg(*Camk2a*-tTA). The strain of origin is FVB. For time-course studies, male and female Tg4510 and WT mice aged 3–8 months were used (at least *n* = 9 per age group). For *in vivo* studies, male and female early-stage (3–3.5 months old) were used. No animal or extracted sample was excluded from any of the analyses. For treatment trials, mice from the same litters were randomly assigned to each experimental group (vehicle n=22 or PACAP n=27) and age-matched WT mice (vehicle n=10 and PACAP n=9). Drug administration and testing were performed in three sets of mice. Mice were housed in 12 h light–12 h dark cycles with free access to food and water.

PS19 mice expressing human TauP301S (1N4R isoform) under control of the mouse prion promoter were obtained from JAX (Stock # 008169, strain name: B6;C3-Tg(Prnp-MAPT^∗^P301S)PS19Vle/J). The colony was maintained as hemizygous Tau PS19 that was used for generating E18 primary neuronal cultures.

### Behavioral testing

#### Morris water maze

The Morris water maze test was carried out as previously described (*125*). After non-spatial training, young (4–4.5 months of age) rTg4510 (27 PACAP and 22 vehicle-treated mice, analyzed in two experiments). A third test was performed on age-matched WT mice (9 PACAP and 10 vehicle - treated). Mice underwent place-discrimination testing for five days with two trials per day. Statistical methods were not employed to predetermine sample size; however, compared to other reported *in vivo* studies with transgenic animals, our studies included higher numbers of animals. Mice from the same litter (4–5 mice per litter) were randomly assigned to each experimental group (vehicle or PACAP). Investigators carrying out tests were blinded to treatment group allocation. No animal was excluded from any of the analyses. We did not apply formal statistical tests for normality or equality of variances but assumed an approximation to the normal distribution when appropriate. Statistical analyses for the Morris water maze test were done by two way repeated measures ANOVA with Bonferroni correction, as described in the statistics section.

#### Novel Object Recognition (NOR) test

NOR testing was performed in a white rectangular open field (60 cm x 50 cm x 26 cm) surrounded by a black curtain. One day before testing, mice were individually habituated to the experimental apparatus without any objects for 5 min. During the training phase, animals were placed in the experimental apparatus with two identical rectangular objects and allowed to explore for 10 min. For the testing phase, one rectangular object was replaced with a cylindrical object and animals were again allowed to explore for 10 min. To test short-term memory, animals were then returned to their cages for a 1 h inter-trial interval. To test long-term memory, animals were returned to their cages for a 24 h inter-trial interval. Before both the training and testing phase, the objects and the apparatus were cleaned with ethanol. All training and testing sessions were recorded and analyzed using automated ANY-maze video tracking software. Exploration of an object was defined as the mouse sniffing the object within 2 cm of it or touching it while facing it. Object placement was balanced to control for potential spatial or side preference. The ability of the mouse to recognize the novel object was determined by dividing the meantime exploring the novel object by the mean of the total time exploring the novel and familiar objects during the test session (T_novel_/[T_novel_ + T_familiar_]. Exploration times were used to calculate a discrimination index (DI) as percent time exploring novel object minus time exploring familiar object divided by total time spent exploring both objects multiplied by 100 ([T_novel_-T_familiar_]/[T_novel_+T_familiar_] x 100). Testers were blind to the treatment group. For statistical analyses, we employed two-way repeated-measures ANOVA with Bonferroni correction described in the statistics section.

### Cell lines

We used the HEK293 (Clone DS1) cell line generated by Dr. Marc Diamond’s lab (*24*) that stably expresses the repeat domain of 2N4R tau with two disease-associated P301L and V337M mutations fused with YFP (RD-P301L/V337M-YFP). These cells allow us to identify tau strains that can seed and template aggregation of endogenous monomeric tau. For seeding assays, DS1 clones were plated on 18mm coverslips coated with 0.05 mg/mL poly-D-lysine and cultured in DMEM containing 10% FBS, 100 U/ml pen /strep and 1% nonessential amino acids. Additionally, we used HEK293 PAC1-EGFP receptor-expressing cell line that was maintained in DMEM/F-12 media containing 10% FBS, 100 U/ml pen /strep, 1% nonessential amino acids.

### Fractionation assay

Postmortem frozen human brains (∼300 mg/sample) or cortices from mice (3 cortices/sample) were homogenized on ice gently with a Heidolph RZR-1 homogenizer in ice-cold homogenization buffer (320 mM sucrose made in hypotonic buffer, 25mM HEPES-KOH pH 7.5, 1mM EDTA, 5mM MES pH 7.5, 5mM MgCl_2_, 5mM ATP pH 7.5, 1mM DTT, 10mM NAF, 25mM beta-glycerol-phosphate, phosphatase, and protease inhibitors). The workflow of the fractionation assay is schematically depicted in Supp. Fig.1A The homogenates were centrifuged at 500 x g for 5 min at 4°C. The supernatant (total extract) was further centrifuged at 19, 000 x g for 20 min at 4°C. The resulting supernatant is a cytosolic fraction, and the pellet is a crude synaptosome that was resuspended in homogenizing buffer.

#### Discontinuous sucrose density gradient

Crude synaptosome extracts were layered on top of non-linear sucrose gradient (1.2M, 0.8M and 0.32M sucrose from bottom to top) and centrifuged at 300, 000 x g for 4 hours at 4°C. Synaptosomes sediment at the interface between 1.2M and 0.8M sucrose layer. To separate synaptosomes into pre and post -synaptic fractions. Collected synaptosomes (∼3–4 mL/sample) were diluted (in 0.01 mM CaCl_2_, 20mM Tris–HCl pH 6, 1% Triton X-100, with protease and phosphatase inhibitors) and mixed by inversion for 20 min at 4°C. After incubation, samples were centrifuged at 40,000 × g for 20 min. The resulting pellet was collected as a post-synaptic fraction, and the supernatant was collected as a pre-synaptic fraction. Following separation of pre and post-synaptic compartments, fractions were precipitated in ice-cold acetone or can be concentrated by Amicon-15 10kDa cut-off tubes. Both fractions were resuspended in PBS with 0.05% Triton X-100 with protease/phosphates inhibitors and sonicated. Fractionation assay was repeated at least three times. Protein concentration was determined by Bradford assay.

#### Continuous Glycerol Density Gradient

Following isolation of synaptosomes, the extracts were subjected to centrifugation at 100,000 × *g* for 24h at 4°C in a 10–40% glycerol gradient. Following ultracentrifugation, 16 fractions (800 μl each) were collected by the fraction collector. Proteins from each fraction were precipitated in ice-cold acetone. Before western blotting, two fractions were combined to one fraction resulting in 8 fractions total that were subjected to western blot analysis.

### Human post-mortem brain tissues

Human tissue samples were provided by the New York Brain Bank at Columbia University Medical Center (CUMC). The demographics of human cases used in this study are listed in **Supplementary Table 1**. The specimens were obtained by consent at autopsy and have been de-identified and are IRB exempt to protect the identity of each patient. At the time of collection, approximately 300 mg block of BA9 region of the brain was dissected out of the frozen brain section and kept at −80°C until homogenization. All cases had been formerly examined by a CUMC neuropathologist to identify a neuropathological diagnosis for the Braak stage.

### In-gel proteasome activity

HEK293 PAC1R-EGFP cells were harvested and homogenized in a buffer containing 50 mM Tris-HCl, pH 7.4, 5 mM MgCl2, 5 mM ATP, 1 mM DTT, phosphatase inhibitors and 10% glycerol, which preserved 26S proteasome assembly and centrifuged at 20,000 × g for 25 min at 4 °C. The supernatant was normalized for protein concentration determined by Bradford assay. Samples (25μg/well lysate) were loaded on a 4% non-denaturing gel and run for 180 min at 160 V. The activity of the 26S proteasome was measured by 100 μM Suc-LLVY-amc (BACHEM Bioscience) diluted in the homogenizing buffer, after incubation for 10-15 minutes at 37^0^C. 26S proteasome bands were detected by trans-illuminator with 365 nm light and photographed by iPhone 10S camera. For immunoblot analysis, native gels were transferred to detect proteasome levels using the 20S proteasome antibody, and phosphorylation status of 26S proteasomes by PKA-specific phospho Ser/ Thre antibody. In-gel assay of 10nM purified 26S proteasomes from *in vivo* study was performed as described here.

### Primary neuronal cultures

Dissociated cortical neurons from E18 PS19 tau of each independent embryo were individually cultured and genotyped to determine Tau PS19 or Non-Tg littermate. Neurons were plated at a density of ∼100,000 cells and mounted on 18 mm glass coverslip (coated with 0.05 mg/mL poly-D-lysine) in neurobasal medium supplemented with 2% B27 and 0.5 mM Glutamax and Pen-Strep. Neurons were cultured for over three weeks in a 37°C incubator with 5% CO_2_. Half of the medium was replaced with fresh medium every 4-5 days. Neurons on day 7-10 *in vitro* were exposed to tau seeds fromDS9 lysate that had been sonicated for 60 pulses just before adding to neurons. Immunocytochemistry was performed at 15-21 d after seeds treatment.

### Intracerebroventricular administration of pumps

Mice were anesthetized with isoflurane for intracerebroventricular administration experiments using Alzet osmotic pumps (model #2004; Alzet Corp, Palo Alto, CA, USA) with a brain infusion kit (Alzet) according to the published protocol (*126*). Osmotic pumps, filled with sterile vehicle (0.9% saline and 0.1% BSA) or PACAP (10 pmol/h; with a rate of 0.25 µl/h) were subcutaneously implanted to allow continuous infusion into the lateral ventricle of the brain. The catheter was placed in the left lateral ventricle with the following coordinates based on bregma: 0.5mm posterior, −1.1 mm lateral (left) and −2.5 mm ventral. Mice were sutured and housed for four weeks. Animals were used in full compliance with the National Institutes of Health/Institutional Animal Care and Use Committee guidelines. The protocol was approved by the Committee on the Ethics of Animal Experiments of Columbia University Medical Center.

### *In Vitro* Seeding Activity of Tau

*In vitro* tau seeding activity was conducted on HEK293 cells and primary neuronal cultures plated on 12-well plates as previously described (*25, 83*).

#### Seeding assay in HEK293 cells

The DS1 clonal line that stably expresses RD-P301L/V337M-YFP were plated in poly-D-lysine 18 mm coverslips at 100,000 cells/well. The next day when cells were at 60–70% confluency and were transduced with 10 µl of lysate normalized to 3ng/µl per well of tau from rTg4510 mice or 5ng/µl per well of tau from human brains plus 1% Lipofectamine 2000 (Invitrogen) diluted in OptiMEM (Thermo Fisher). Cells were incubated with lysate from rTg4510 or human brains for 24 or 48 h, respectively, before fixation with 4% PFA for immunofluorescence (see below).

#### Seeding assay in primary neuronal cultures

Primary cortical neurons from PS19 mice were plated at density 100, 000 and were exposed at 14 days in vitro (DIV) to 25ug lysates from HEK293-DS9 clone that feature tau aggregates and have high seeding activity. DS9 lysate was used as tau seeds to induce aggregation and propagation of tau in naïve neuronal cells. Neurons were exposed to seeds for 5-7 days. To test whether PACAP can attenuate seeding activity, neurons were treated for four days before they were fixed for immunocytochemistry experiments. During fixation with 4% PFA, 0.1% Triton X-100 was used to remove soluble tau, and primary neurons were stained for conformationally altered tau (MC1) 5-7 days following seeding (Fig 3). In other experimental paradigms, we omitted Triton X-100 to assess neurite integrity during seed exposure (Supplementary Fig. 4).

### Immunoblot analysis

Samples (5–10 µg protein) were typically run on 4–12% Bis-Tris gels (Life Technologies; WG1403BOX10) using MOPS buffer (NP0001) with antioxidant (NP0005). Proteins were analyzed after electrophoresis on SDS-PAGE and transferred onto 0.2-µm nitrocellulose membranes (Whatman). Blots were blocked and incubated with primary and secondary antibodies at concentrations listed below. Membranes were developed with enhanced chemiluminescent reagent (Immobilon Western HRP substrate and Luminol reagent (WBKLS0500, Millipore) using a Fujifilm LAS3000 imaging system. ImageJ (http://rsb.info.nih.gov/ij) was used to quantify the signal. Relative intensity (fold change or fold increase, no units) is the ratio of the value for each protein to the value of the respective loading control.

### Immunofluorescence

#### Primary neuronal cells and HEK293 (DS1) cells

Following treatment, neurons and DS1 cells were rinsed three times in PBS and fixed with 4% paraformaldehyde for 15 min at RT. In the case of neurons, 0.1% Triton X-100 was used to remove soluble tau, to allow for the detection of insoluble aggregates that stain positive for conformationally altered tau (MC1) 7days the following seeding. Alternatively, seeded neuronal cultures were fixed without Triton X-100. Afterward, cells were blocked for 1 h in 5% bovine serum albumin/PBS and were processed for immunofluorescence using the following primary antibodies: mouse anti-MC1 (1:500) and rabbit anti-MAP2 (1:500) for 24 h at 4 °C. Fluorescent-conjugated secondary antisera mixtures containing Alexa 488 IgG and Alexa 594 IgG (1:500) (anti-mouse and anti-rabbit Alexa, Invitrogen Molecular Probes) were used, respectively. Cells were mounted on coverslips with Prolong Gold antifade containing DAPI (Invitrogen). In the case of HEK293 cells after fixation, no antibody was used; instead, cells were mounted on coverslips for confocal imaging.

#### Mouse brain tissue

Brains of rTg4510 mice were isolated after transcranial perfusion with PBS, and one hemisphere was drop-fixed in 4% PFA overnight then subject to cryoprotection treatment in 30% sucrose in PBS for 24 h. Free-floating brain sections (35 µm) from brains sectioned in the sagittal plane were used. The sections were incubated at 4 °C overnight with primary antibody diluted in PBS containing 0.3% Triton X-100 and 5% normal goat serum blocking solution (Vector Laboratories, #S-1000). Antibodies were as follows: anti-mouse monoclonal pS396 and pS404 (PHF1-1:1,000). Following washes, sections were incubated with goat anti-mouse IgG Alexa 594 (1:500).

Staining was visualized by confocal microscopy, Zeiss LSM710 confocal microscope at 20× dry and 40× oil immersion objectives. Sequential tile scans were performed to capture images of 1,024 × 1,024 resolution. All images from the same experiment were taken at the same laser intensity and detector gain.

### Purification of 26S proteasomes

Cortices from 6 mice were pooled each time proteasome purification was carried out, and 26S proteasomes were affinity purified using a UBL domain as the ligand. Briefly, brain homogenates were spun for 1 h at 100,000 × *g*. The soluble extracts were incubated at 4 °C with 2 mg/ml glutathione-*S*-transferase–ubiquitin-like domain (GST-UBL) and 300 µL of glutathione–Sepharose 4B (GSH-Sepharose). The slurry containing 26S proteasomes bound to GST-UBL was poured into an empty column and washed, then incubated with 2 mg/ml His_10_-ubiquitin–interacting motif (10× His-UIM). The eluate was collected and incubated with Ni^2+^-NTA-agarose for 20 min at 4 °C. The Ni^2+^-NTA-bound 10× His-UIM was removed by filtration. The resulting flow-through (∼0.6 mL) contained purified 26S proteasomes. The molarity of 26S proteasome particles was calculated, assuming a molecular weight of 2.5 MDa. To assess the peptidase capacity of 26S proteasome subunits, we incubated 10 nM of the proteasome with 40 µM Suc-LLVY-amc fluorogenic peptide for chymotrypsin-like activity (β5 activity). Kinetic reactions (peptidase activity) were carried out for a period of 60-120 min. The rate of the degradation of the substrate over time was calculated as the slope of the reaction. The fluorescence signal was captured at 380 nm excitation, 460 nm emission by Infinite 200 PRO multimode reader (TECAN).

### Tissue fractionation and protein extraction for western blotting

Mice were sacrificed by cervical dislocation, and the brains were immediately dissected on wet ice and stored on dry ice. Briefly, frozen hemispheres free of cerebellum and brainstem were weighed and homogenized without thawing in RIPA buffer (10× volume/weight) (50 mM Tris-HCl, pH 7.4, 1% NP-40, 0.25% sodium deoxycholate, 150 mM NaCl, 1 mM EDTA, 1 mM phenylmethyl-sulfonyl (PMSF), 1 mM sodium orthovanadate, 1 mM sodium fluoride (NaF), 1 µl/ml protease inhibitor mix (Sigma-Aldrich)). Homogenates were centrifuged for 10 min at 3,000 × *g* at 4 °C. Protein assay was performed on the clear supernatants representing the total extract used for the analysis of the total protein levels. Sample volumes were adjusted with RIPA buffer containing 100 mMDTT and NuPAGE LDS Sample Buffer 4× buffer (Life Technologies) and boiled for 5 min. The sarkosyl-insoluble extracts, which are highly enriched in aggregated tau species, were generated when 200 µg aliquots from the total protein extracts were normalized into 200 µl final volume containing 1% sarkosyl, followed by ultra-centrifugation at 100,000 × *g* for 1 h at 20 °C. Without disturbing the pellet, the supernatant was transferred to new tubes. The pellet was resuspended in 100 µL RIPA buffer containing DTT and NuPAGE LDS Sample Buffer 4× buffer, followed by vortexing for 1 min and 5 min heating at 95 °C. The heat-stable extract, which contains soluble tau, was obtained when the supernatant was further processed, first by heating for 5 min at 95 °C followed by 30 min centrifugation at 20,000 × *g*. Extracts were transferred to new tubes containing NuPAGE LDS Sample Buffer 4× buffer (4:1 (extracts/buffer) ratio).

### Tau ELISA

Sandwich ELISA was performed as previously described using tau monoclonal antibodies DA31 pan-tau antibody that maps in the amino acid region 150–190 of tau and DA9-HRP (gifts from Dr. Peter Davies) (*127*). Briefly, 96-well plates were coated with the DA31 antibody at a final concentration of 6 µg/ml in coating buffer for at least 48 h at 4°C. After washing 3X in the wash buffer, the plates were blocked for 1 h at room temperature using StartingBlock (Thermo scientific) to avoid non-specific binding. Each plate was then washed 5 times and 50 µl of diluted samples were added to the wells. Concurrently, 50 µl of the DA9-HRP detection antibody (dilution 1/50) was added to the samples. Plates were incubated O/N shaking at 4°C and then washed 5 times in wash buffer. 1-Step ULTRA TMB-ELISA (Thermo scientific) was added for 30 minutes at room temperature before stopping the reaction with 2 M H_2_SO_4_. Plates were read with an Infinite m200 plate reader (Tecan) at 450 nm.

### Statistical analyses

Statistical analyses were performed with Prism6 (Graphpad Software, San Diego, CA). Data from fractionation assays with WT and rTg4510 mice, *in vitro* seeding assays with DS1-HEK293 cells, Morris water maze, and novel object recognition tests, utilized two-way repeated-measures ANOVA with post hoc Bonferroni correction. For fractionation assays with human samples, and with rTg4510 mice from *in vivo* studies, and *in vitro* seeding assay in primary neuronal cultures and proteasome assays, we employed one-way ANOVA followed by Bonferroni multiple comparison *post hoc* test. Data from immunoblot analysis from the total and insoluble extracts and discrimination index from novel object recognition and from purified extracts of proteasome for PKA-specific phospho Ser/Thre were analyzed using unpaired two-tailed Student’s *t*-test between groups with unequal variance.

---

**Supplementary Figure 1.**
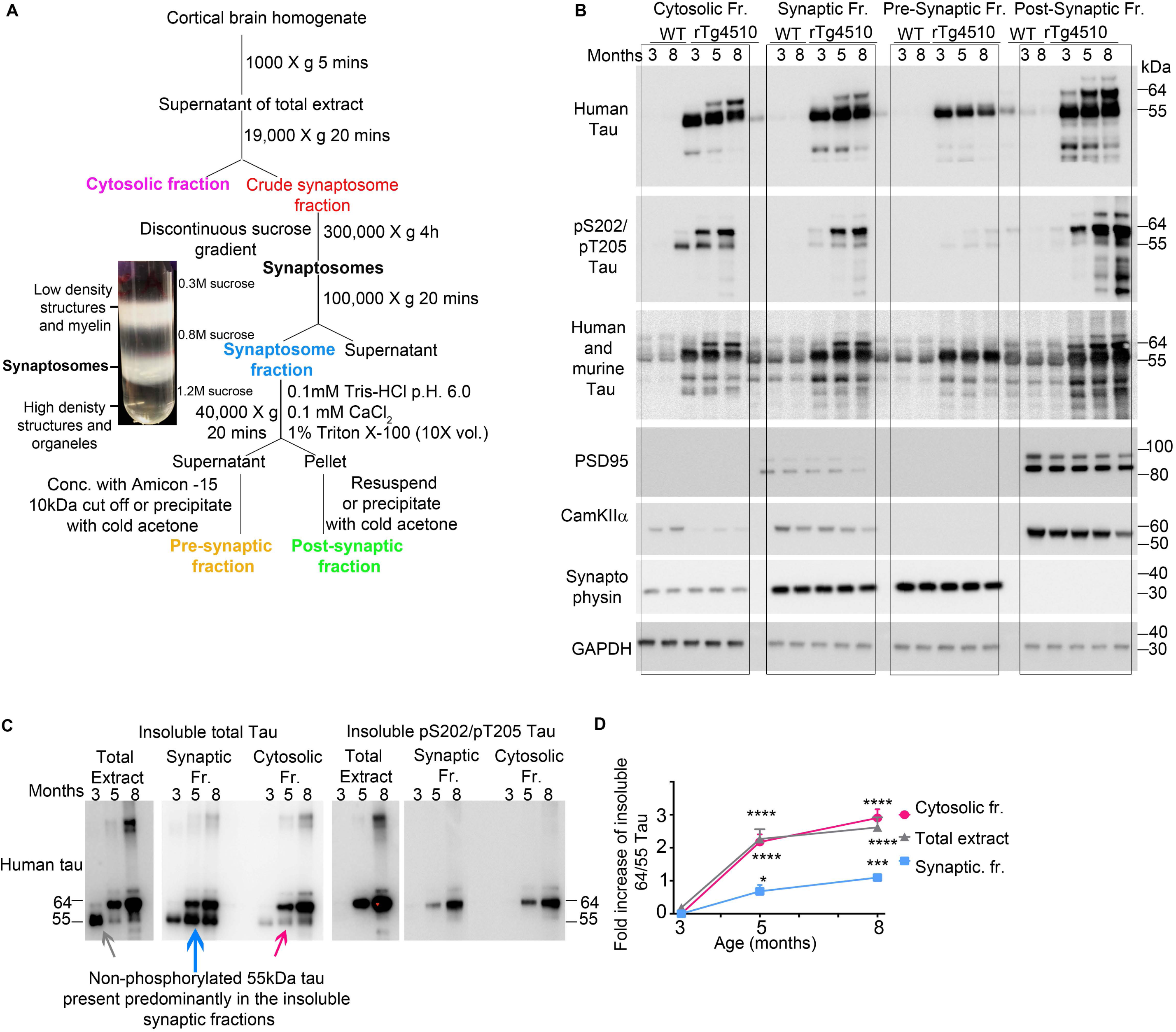
(**A**) Schematic diagram of the subcellular fractionation protocol to generate cytosolic, synaptic, pre-synaptic and post-synaptic fractions from post-mortem humans and mice tissue. (**B**) Representative uncut immunoblotting data from Fig 1A. (**C**) A representative of immunoblot analysis of insoluble extracts from the total, cytosolic and synaptic fractions for tau and pS202 and pS205 tau epitope. (**D**) The quantitative analysis represented as a fold increase of insoluble 64/55 tau ratio relative to 3 months of age. Two-way repeated-measures ANOVA with post hoc Bonferroni correction was employed. Error bars, mean ± SEM. ***P < 0.001, ****P<.0.0001.

**Supplementary Figure 2.**
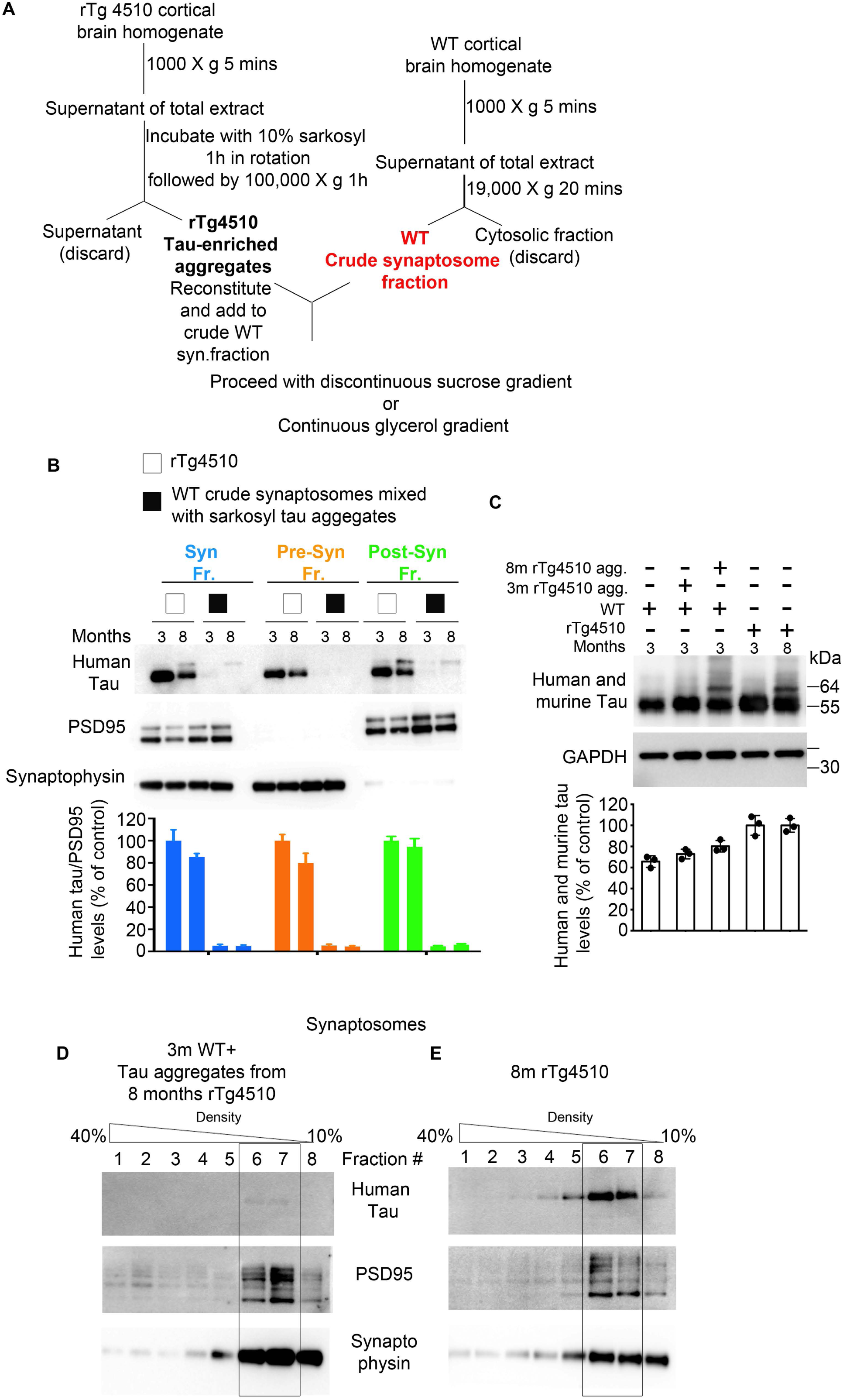
(**A**) Schematic diagram of the subcellular fractionation protocol from crude synaptosomes of WT mice mixed with sarkosyl tau aggregates to generate synaptic, pre and post -synaptic fractions. (**B**) The representative of immunoblot analysis for synaptic, pre and post -synaptic fractions of rTg4510 and WT mice mixed with sarkosyl tau aggregates from 3 and 8 months old rTg4510 mice. (**C**) Total extracts before the fractionation procedure. Continuous Glycerol Density Gradient (10-40% density) of isolated synaptosomes from: (**D**) 3 months old WT crude synaptosomes mixed with tau aggregates from 8months old rTg4510 and (**E**) synaptosomes from 8 months rTg4510.

**Supplementary Figure 3.**
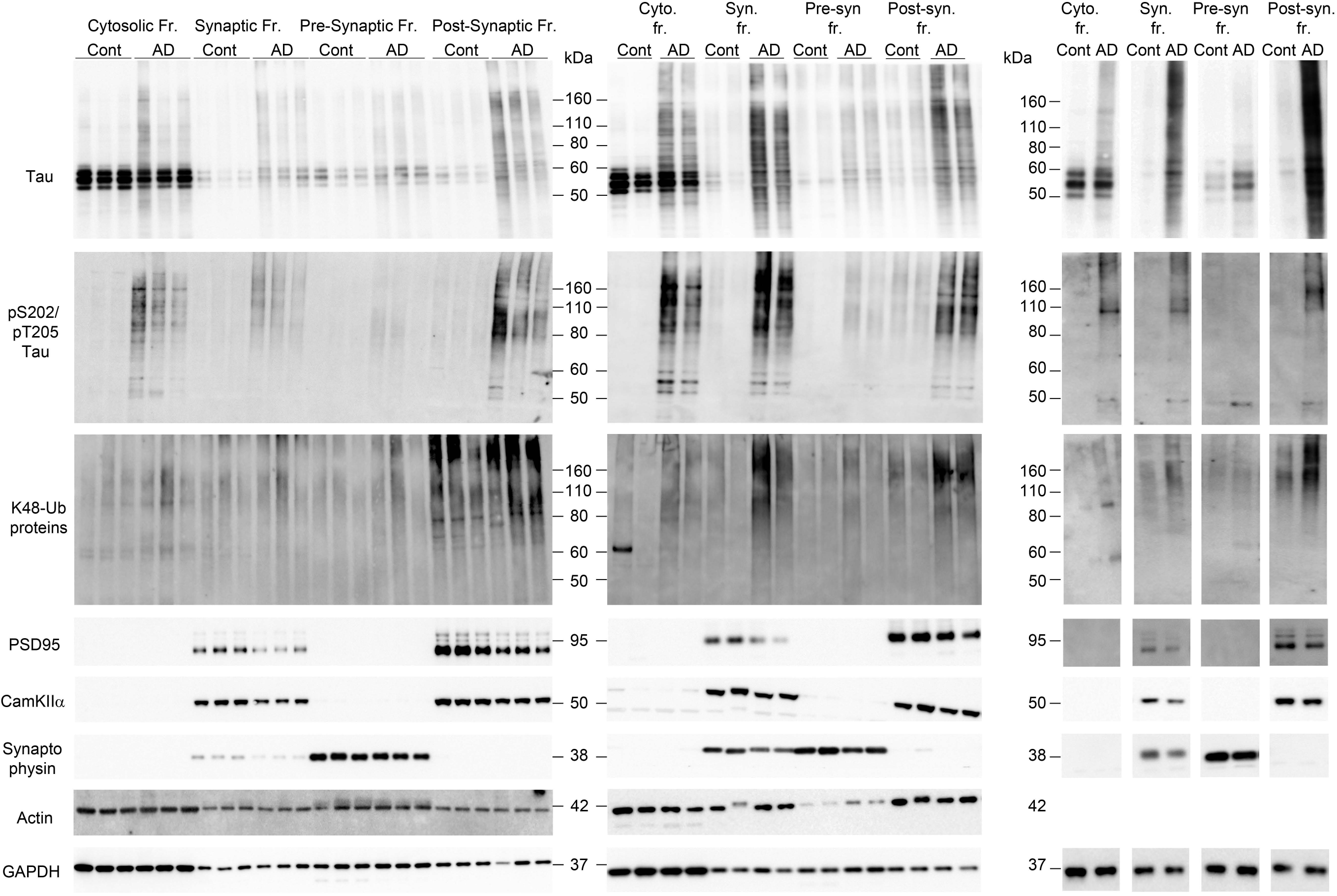
Summary of uncut immunoblotting data from human post mortem subcellular fractionation assay from Fig. 1 D.

**Supplementary Figure 4.**
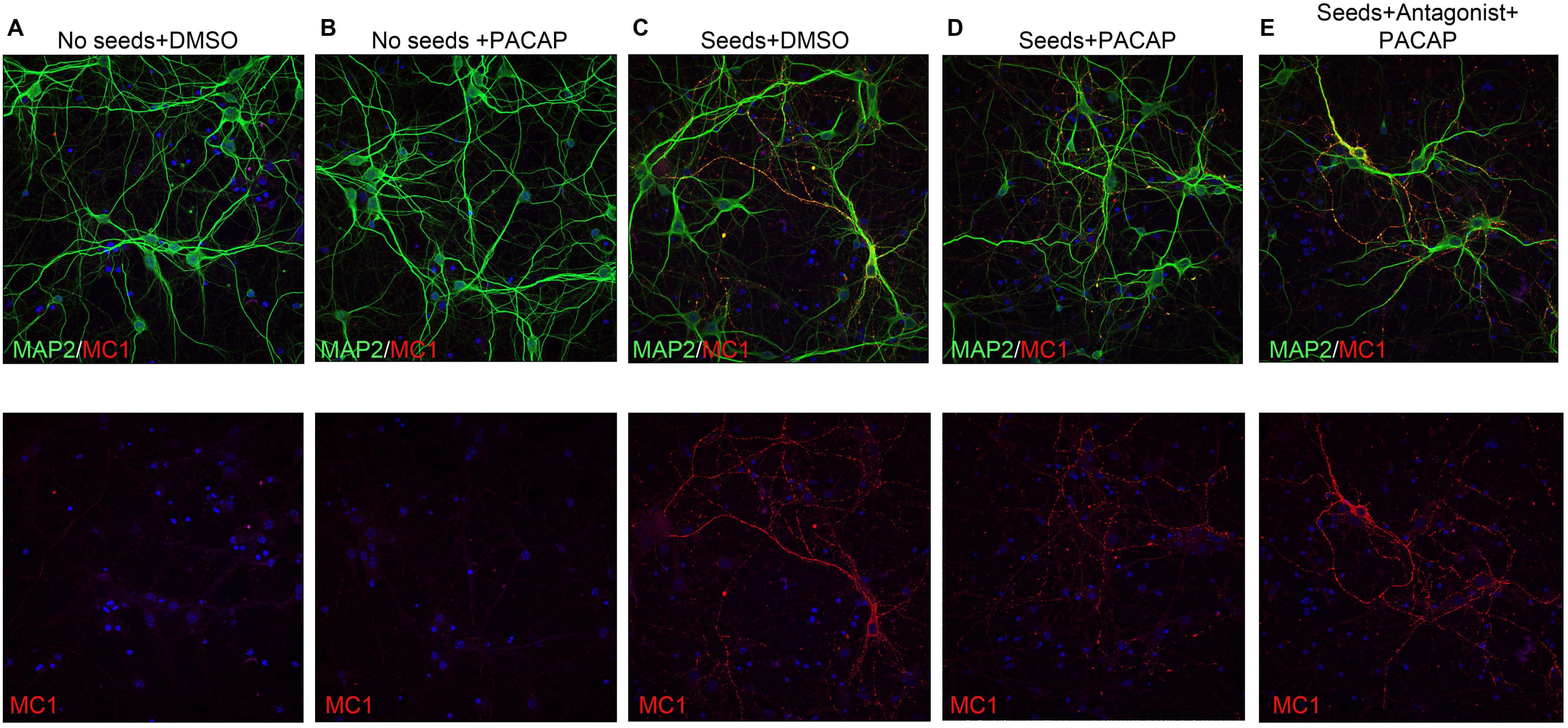
Representative images for MC1 and MAP2 (merged) or MC1 alone of PS19 of primary neuronal cultures treated with (**A**) DMSO or (**B**) PACAP for four days followed by fluorescent immunostaining. Representative images of PS19 of primary neuronal cultures (**C, D, E**) seeded with 20 µg of DS9 cell lysate followed by four days treatment with (**C**) DMSO, (**D**) 100 nM PACAP or (**E**) pre-treated with 200nM PAC1R antagonist for 6 hours before adding 100 nM PACAP. Cells were fixed 5-7 days post-seed exposure without 0.1% Triton X-100 and immunostained for MC1 and MAP2.

**Supplementary Figure 5.**
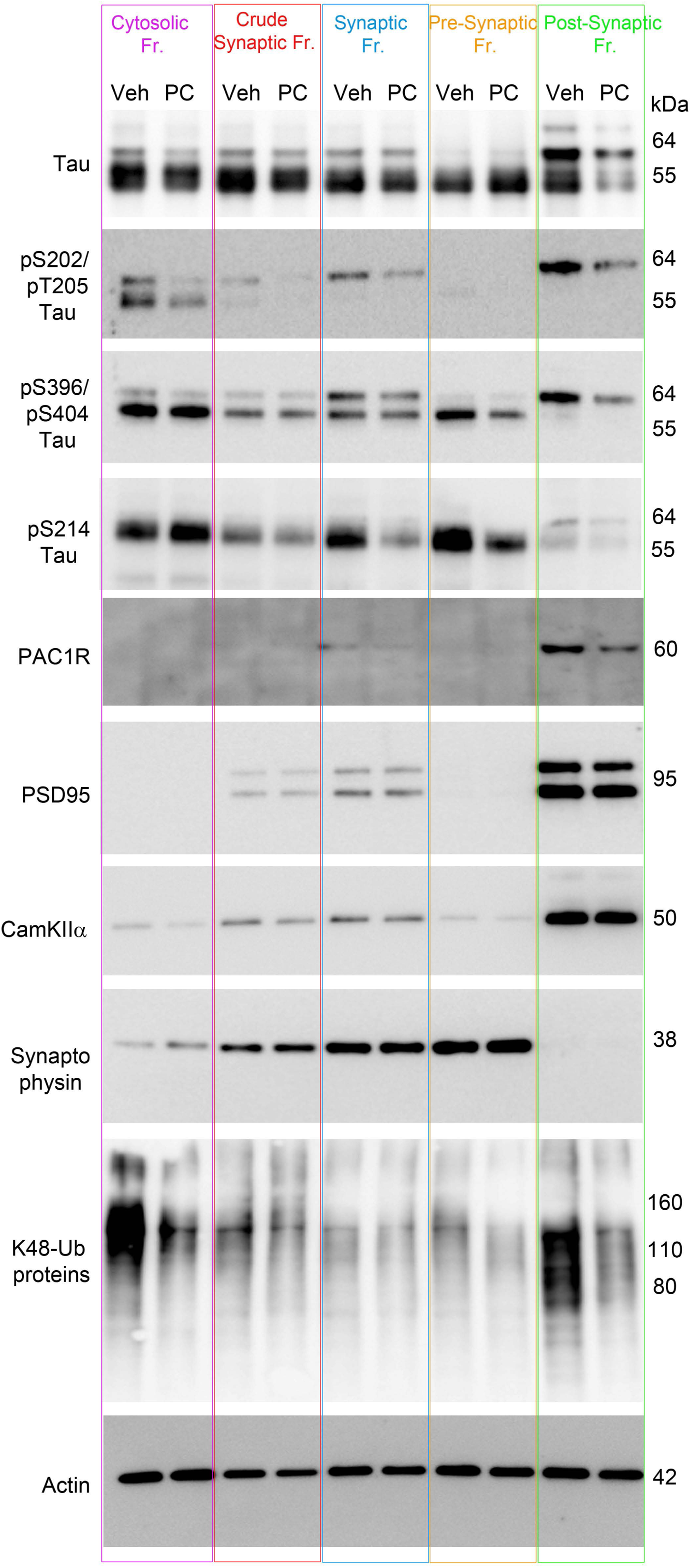
The representative of uncut immunoblotting data from Fig. 4.

**Supplementary Figure 6.**
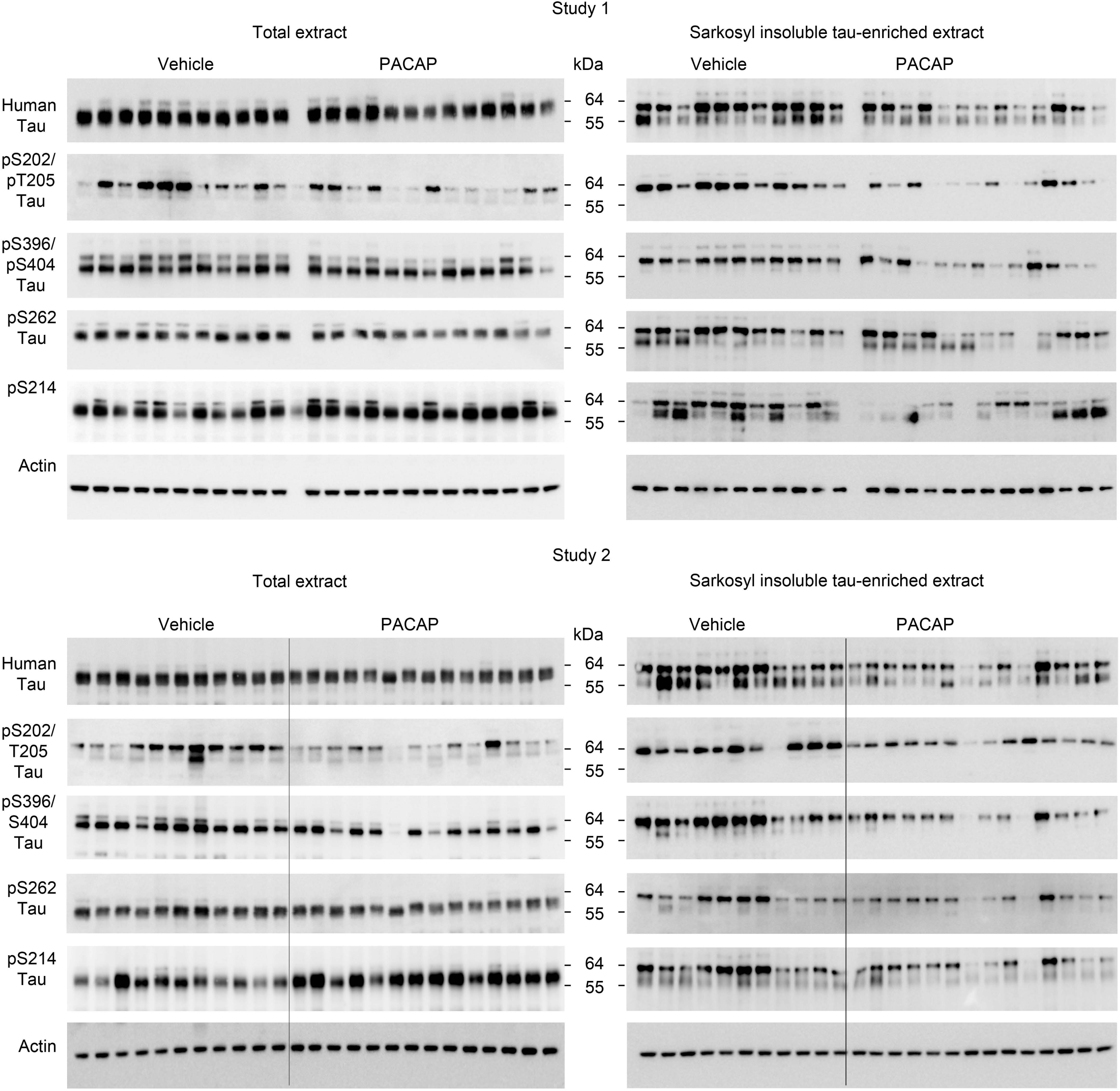
Summary of uncut immunoblotting data from Fig.6. *In vivo* treatments were carried out in two sets.

**TABLE 1:**
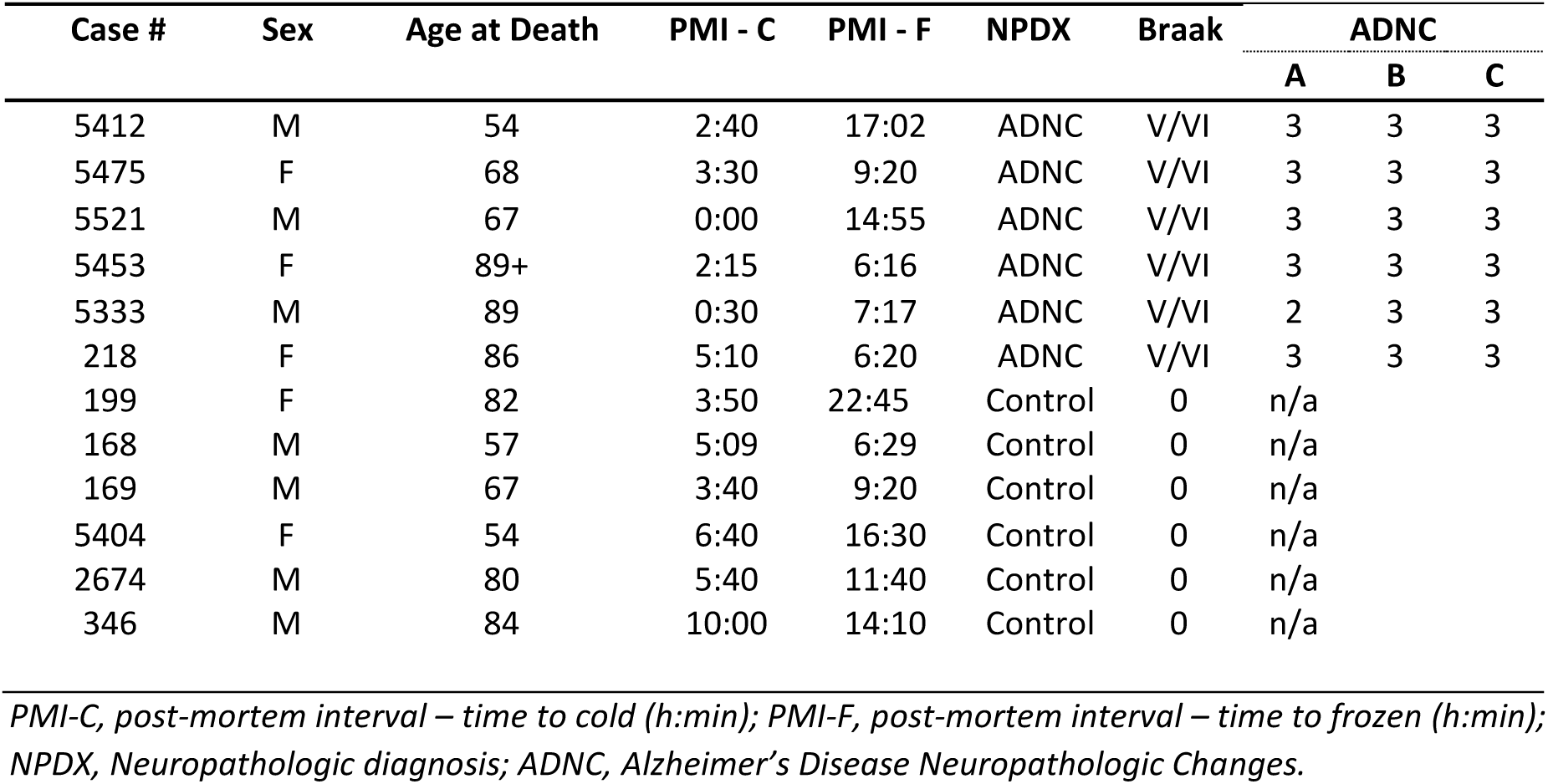
Human demographics and neuropathological data.

| Case # | Sex | Age at Death | PMI - C | PMI - F | NPDX | Braak | ADNC |  |  |
| --- | --- | --- | --- | --- | --- | --- | --- | --- | --- |
|  |  |  |  |  |  |  | A | B | C |
| 5412 | M | 54 | 2:40 | 17:02 | ADNC | V/VI | 3 | 3 | 3 |
| 5475 | F | 68 | 3:30 | 9:20 | ADNC | V/VI | 3 | 3 | 3 |
| 5521 | M | 67 | 0:00 | 14:55 | ADNC | V/VI | 3 | 3 | 3 |
| 5453 | F | 89+ | 2:15 | 6:16 | ADNC | V/VI | 3 | 3 | 3 |
| 5333 | M | 89 | 0:30 | 7:17 | ADNC | V/VI | 2 | 3 | 3 |
| 218 | F | 86 | 5:10 | 6:20 | ADNC | V/VI | 3 | 3 | 3 |
| 199 | F | 82 | 3:50 | 22:45 | Control | 0 | n/a |  |  |
| 168 | M | 57 | 5:09 | 6:29 | Control | 0 | n/a |  |  |
| 169 | M | 67 | 3:40 | 9:20 | Control | 0 | n/a |  |  |
| 5404 | F | 54 | 6:40 | 16:30 | Control | 0 | n/a |  |  |
| 2674 | M | 80 | 5:40 | 11:40 | Control | 0 | n/a |  |  |
| 346 | M | 84 | 10:00 | 14:10 | Control | 0 | n/a |  |  |

*PMI-C, post-mortem interval – time to cold (h:min); PMI-F, post-mortem interval – time to frozen (h:min); NPDX, Neuropathologic diagnosis; ADNC, Alzheimer's Disease Neuropathologic Changes.*

## References

1. C. A. Davies, D. M. Mann, P. Q. Sumpter, P. O. Yates, A quantitative morphometric analysis of the neuronal and synaptic content of the frontal and temporal cortex in patients with Alzheimer’s disease. J Neurol Sci 78, 151–164 (1987); published online EpubApr (

2. E. Masliah, M. Mallory, M. Alford, R. DeTeresa, L. A. Hansen, D. W. McKeel, Jr., J. C. Morris, Altered expression of synaptic proteins occurs early during progression of Alzheimer’s disease. Neurology 56, 127–129 (2001); published online EpubJan 09 (

3. S. W. Scheff, D. A. Price, F. A. Schmitt, S. T. DeKosky, E. J. Mufson, Synaptic alterations in CA1 in mild Alzheimer disease and mild cognitive impairment. Neurology 68, 1501–1508 (2007); published online EpubMay 01 (10.1212/01.wnl.0000260698.46517.8f).

4. M. K. Chen, A. P. Mecca, M. Naganawa, S. J. Finnema, T. Toyonaga, S. F. Lin, S. Najafzadeh, J. Ropchan, Y. Lu, J. W. McDonald, H. R. Michalak, N. B. Nabulsi, A. F. T. Arnsten, Y. Huang, R. E. Carson, C. H. van Dyck, Assessing Synaptic Density in Alzheimer Disease With Synaptic Vesicle Glycoprotein 2A Positron Emission Tomographic Imaging. JAMA Neurol 75, 1215–1224 (2018); published online EpubOct 1 (10.1001/jamaneurol.2018.1836).

5. S. T. DeKosky, S. W. Scheff, Synapse loss in frontal cortex biopsies in Alzheimer’s disease: correlation with cognitive severity. Ann Neurol 27, 457–464 (1990); published online EpubMay (10.1002/ana.410270502).

6. A. Bejanin, D. R. Schonhaut, R. La Joie, J. H. Kramer, S. L. Baker, N. Sosa, N. Ayakta, A. Cantwell, M. Janabi, M. Lauriola, J. P. O’Neil, M. L. Gorno-Tempini, Z. A. Miller, H. J. Rosen, B. L. Miller, W. J. Jagust, G. D. Rabinovici, Tau pathology and neurodegeneration contribute to cognitive impairment in Alzheimer’s disease. Brain : a journal of neurology 140, 3286–3300 (2017); published online EpubDec 1 (10.1093/brain/awx243).

7. P. V. Arriagada, K. Marzloff, B. T. Hyman, Distribution of Alzheimer-type pathologic changes in nondemented elderly individuals matches the pattern in Alzheimer’s disease. Neurology 42, 1681–1688 (1992); published online EpubSep (

8. J. Zhou, E. D. Gennatas, J. H. Kramer, B. L. Miller, W. W. Seeley, Predicting regional neurodegeneration from the healthy brain functional connectome. Neuron 73, 1216–1227 (2012); published online EpubMar 22 (10.1016/j.neuron.2012.03.004).

9. C. Xia, S. J. Makaretz, C. Caso, S. McGinnis, S. N. Gomperts, J. Sepulcre, T. Gomez-Isla, B. T. Hyman, A. Schultz, N. Vasdev, K. A. Johnson, B. C. Dickerson, Association of In Vivo [18F]AV-1451 Tau PET Imaging Results With Cortical Atrophy and Symptoms in Typical and Atypical Alzheimer Disease. JAMA neurology 74, 427–436 (2017); published online EpubApr 1 (10.1001/jamaneurol.2016.5755).

10. B. G. Perez-Nievas, T. D. Stein, H. C. Tai, O. Dols-Icardo, T. C. Scotton, I. Barroeta-Espar, L. Fernandez-Carballo, E. L. de Munain, J. Perez, M. Marquie, A. Serrano-Pozo, M. P. Frosch, V. Lowe, J. E. Parisi, R. C. Petersen, M. D. Ikonomovic, O. L. Lopez, W. Klunk, B. T. Hyman, T. Gomez-Isla, Dissecting phenotypic traits linked to human resilience to Alzheimer’s pathology. Brain 136, 2510–2526 (2013); published online EpubAug (10.1093/brain/awt171).

11. H. C. Tai, B. Y. Wang, A. Serrano-Pozo, M. P. Frosch, T. L. Spires-Jones, B. T. Hyman, Frequent and symmetric deposition of misfolded tau oligomers within presynaptic and postsynaptic terminals in Alzheimer’s disease. Acta neuropathologica communications 2, 146 (2014); published online EpubOct 21 (10.1186/s40478-014-0146-2).

12. E. Braak, H. Braak, E. M. Mandelkow, A sequence of cytoskeleton changes related to the formation of neurofibrillary tangles and neuropil threads. Acta neuropathologica 87, 554–567 (1994).

13. H. Braak, D. R. Thal, E. Ghebremedhin, K. Del Tredici, Stages of the pathologic process in Alzheimer disease: age categories from 1 to 100 years. Journal of neuropathology and experimental neurology 70, 960–969 (2011); published online EpubNov (10.1097/NEN.0b013e318232a379).

14. B. R. Hoover, M. N. Reed, J. Su, R. D. Penrod, L. A. Kotilinek, M. K. Grant, R. Pitstick, G. A. Carlson, L. M. Lanier, L. L. Yuan, K. H. Ashe, D. Liao, Tau mislocalization to dendritic spines mediates synaptic dysfunction independently of neurodegeneration. Neuron 68, 1067–1081 (2010); published online EpubDec 22 (10.1016/j.neuron.2010.11.030).

15. V. Balaji, S. Kaniyappan, E. Mandelkow, Y. Wang, E. M. Mandelkow, Pathological missorting of endogenous MAPT/Tau in neurons caused by failure of protein degradation systems. Autophagy 14, 2139–2154 (2018)10.1080/15548627.2018.1509607).

16. D. Xia, C. Li, J. Gotz, Pseudophosphorylation of Tau at distinct epitopes or the presence of the P301L mutation targets the microtubule-associated protein Tau to dendritic spines. Biochimica et biophysica acta 1852, 913–924 (2015); published online EpubMay (10.1016/j.bbadis.2014.12.017).

17. T. E. Cope, T. Rittman, R. J. Borchert, P. S. Jones, D. Vatansever, K. Allinson, L. Passamonti, P. Vazquez Rodriguez, W. R. Bevan-Jones, J. T. O’Brien, J. B. Rowe, Tau burden and the functional connectome in Alzheimer’s disease and progressive supranuclear palsy. Brain : a journal of neurology 141, 550–567 (2018); published online EpubFeb 1 (10.1093/brain/awx347).

18. L. Liu, V. Drouet, J. W. Wu, M. P. Witter, S. A. Small, C. Clelland, K. Duff, Trans-synaptic spread of tau pathology in vivo. PloS one 7, e31302 (2012)10.1371/journal.pone.0031302).

19. A. de Calignon, M. Polydoro, M. Suarez-Calvet, C. William, D. H. Adamowicz, K. J. Kopeikina, R. Pitstick, N. Sahara, K. H. Ashe, G. A. Carlson, T. L. Spires-Jones, B. T. Hyman, Propagation of tau pathology in a model of early Alzheimer’s disease. Neuron 73, 685–697 (2012); published online EpubFeb 23 (10.1016/j.neuron.2011.11.033).

20. I. C. Stancu, B. Vasconcelos, L. Ris, P. Wang, A. Villers, E. Peeraer, A. Buist, D. Terwel, P. Baatsen, T. Oyelami, N. Pierrot, C. Casteels, G. Bormans, P. Kienlen-Campard, J. N. Octave, D. Moechars, I. Dewachter, Templated misfolding of Tau by prion-like seeding along neuronal connections impairs neuronal network function and associated behavioral outcomes in Tau transgenic mice. Acta neuropathologica 129, 875–894 (2015); published online EpubJun (10.1007/s00401-015-1413-4).

21. F. Clavaguera, T. Bolmont, R. A. Crowther, D. Abramowski, S. Frank, A. Probst, G. Fraser, A. K. Stalder, M. Beibel, M. Staufenbiel, M. Jucker, M. Goedert, M. Tolnay, Transmission and spreading of tauopathy in transgenic mouse brain. Nature cell biology 11, 909–913 (2009); published online EpubJul (10.1038/ncb1901).

22. M. Iba, J. D. McBride, J. L. Guo, B. Zhang, J. Q. Trojanowski, V. M. Lee, Tau pathology spread in PS19 tau transgenic mice following locus coeruleus (LC) injections of synthetic tau fibrils is determined by the LC’s afferent and efferent connections. Acta neuropathologica 130, 349–362 (2015); published online EpubSep (10.1007/s00401-015-1458-4).

23. E. Peeraer, A. Bottelbergs, K. Van Kolen, I. C. Stancu, B. Vasconcelos, M. Mahieu, H. Duytschaever, L. Ver Donck, A. Torremans, E. Sluydts, N. Van Acker, J. A. Kemp, M. Mercken, K. R. Brunden, J. Q. Trojanowski, I. Dewachter, V. M. Lee, D. Moechars, Intracerebral injection of preformed synthetic tau fibrils initiates widespread tauopathy and neuronal loss in the brains of tau transgenic mice. Neurobiology of disease 73, 83–95 (2015); published online EpubJan (10.1016/j.nbd.2014.08.032).

24. D. W. Sanders, S. K. Kaufman, S. L. DeVos, A. M. Sharma, H. Mirbaha, A. Li, S. J. Barker, A. C. Foley, J. R. Thorpe, L. C. Serpell, T. M. Miller, L. T. Grinberg, W. W. Seeley, M. I. Diamond, Distinct tau prion strains propagate in cells and mice and define different tauopathies. Neuron 82, 1271–1288 (2014); published online EpubJun 18 (10.1016/j.neuron.2014.04.047).

25. B. B. Holmes, J. L. Furman, T. E. Mahan, T. R. Yamasaki, H. Mirbaha, W. C. Eades, L. Belaygorod, N. J. Cairns, D. M. Holtzman, M. I. Diamond, Proteopathic tau seeding predicts tauopathy in vivo. Proceedings of the National Academy of Sciences of the United States of America 111, E4376–4385 (2014); published online EpubOct 14 (10.1073/pnas.1411649111).

26. S. Boluda, M. Iba, B. Zhang, K. M. Raible, V. M. Lee, J. Q. Trojanowski, Differential induction and spread of tau pathology in young PS19 tau transgenic mice following intracerebral injections of pathological tau from Alzheimer’s disease or corticobasal degeneration brains. Acta neuropathologica 129, 221–237 (2015); published online EpubFeb (10.1007/s00401-014-1373-0).

27. F. Clavaguera, H. Akatsu, G. Fraser, R. A. Crowther, S. Frank, J. Hench, A. Probst, D. T. Winkler, J. Reichwald, M. Staufenbiel, B. Ghetti, M. Goedert, M. Tolnay, Brain homogenates from human tauopathies induce tau inclusions in mouse brain. Proceedings of the National Academy of Sciences of the United States of America 110, 9535–9540 (2013); published online EpubJun 4 (10.1073/pnas.1301175110).

28. H. Braak, E. Braak, Neuropathological stageing of Alzheimer-related changes. Acta neuropathologica 82, 239–259 (1991).

29. H. C. Tai, A. Serrano-Pozo, T. Hashimoto, M. P. Frosch, T. L. Spires-Jones, B. T. Hyman, The synaptic accumulation of hyperphosphorylated tau oligomers in Alzheimer disease is associated with dysfunction of the ubiquitin-proteasome system. The American journal of pathology 181, 1426–1435 (2012); published online EpubOct (10.1016/j.ajpath.2012.06.033).

30. B. Bingol, E. M. Schuman, Synaptic protein degradation by the ubiquitin proteasome system. Current opinion in neurobiology 15, 536–541 (2005); published online EpubOct (10.1016/j.conb.2005.08.016).

31. L. D. Cohen, N. E. Ziv, Recent insights on principles of synaptic protein degradation. F1000Research 6, 675 (2017)10.12688/f1000research.10599.1).

32. S. tom Dieck, L. Kochen, C. Hanus, M. Heumuller, I. Bartnik, B. Nassim-Assir, K. Merk, T. Mosler, S. Garg, S. Bunse, D. A. Tirrell, E. M. Schuman, Direct visualization of newly synthesized target proteins in situ. Nature methods 12, 411–414 (2015); published online EpubMay (10.1038/nmeth.3319).

33. A. N. Hegde, Ubiquitin-proteasome-mediated local protein degradation and synaptic plasticity. Progress in neurobiology 73, 311–357 (2004); published online EpubAug (10.1016/j.pneurobio.2004.05.005).

34. A. N. Hegde, Proteolysis, synaptic plasticity and memory. Neurobiology of learning and memory 138, 98–110 (2017); published online EpubFeb (10.1016/j.nlm.2016.09.003).

35. J. Felsenberg, V. Dombrowski, D. Eisenhardt, A role of protein degradation in memory consolidation after initial learning and extinction learning in the honeybee (Apis mellifera). Learning & memory 19, 470–477 (2012); published online EpubSep 17 (10.1101/lm.026245.112).

36. B. K. Kaang, J. H. Choi, Synaptic protein degradation in memory reorganization. Advances in experimental medicine and biology 970, 221–240 (2012)10.1007/978-3-7091-0932-8_10).

37. T. J. Jarome, F. J. Helmstetter, The ubiquitin-proteasome system as a critical regulator of synaptic plasticity and long-term memory formation. Neurobiology of learning and memory 105, 107–116 (2013); published online EpubOct (10.1016/j.nlm.2013.03.009).

38. G. A. Collins, A. L. Goldberg, The Logic of the 26S Proteasome. Cell 169, 792–806 (2017); published online EpubMay 18 (10.1016/j.cell.2017.04.023).

39. J. N. Keller, K. B. Hanni, W. R. Markesbery, Impaired proteasome function in Alzheimer’s disease. Journal of neurochemistry 75, 436–439 (2000); published online EpubJul (

40. N. Myeku, C. L. Clelland, S. Emrani, N. V. Kukushkin, W. H. Yu, A. L. Goldberg, K. E. Duff, Tau-driven 26S proteasome impairment and cognitive dysfunction can be prevented early in disease by activating cAMP-PKA signaling. Nat Med 22, 46–53 (2016); published online EpubJan (10.1038/nm.4011).

41. P. Deriziotis, R. Andre, D. M. Smith, R. Goold, K. J. Kinghorn, M. Kristiansen, J. A. Nathan, R. Rosenzweig, D. Krutauz, M. H. Glickman, J. Collinge, A. L. Goldberg, S. J. Tabrizi, Misfolded PrP impairs the UPS by interaction with the 20S proteasome and inhibition of substrate entry. The EMBO journal 30, 3065–3077 (2011); published online EpubAug 3 (10.1038/emboj.2011.224).

42. M. Kristiansen, P. Deriziotis, D. E. Dimcheff, G. S. Jackson, H. Ovaa, H. Naumann, A. R. Clarke, F. W. van Leeuwen, V. Menendez-Benito, N. P. Dantuma, J. L. Portis, J. Collinge, S. J. Tabrizi, Disease-associated prion protein oligomers inhibit the 26S proteasome. Molecular cell 26, 175–188 (2007); published online EpubApr 27 (10.1016/j.molcel.2007.04.001).

43. R. Andre, S. J. Tabrizi, Misfolded PrP and a novel mechanism of proteasome inhibition. Prion 6, 32–36 (2012); published online EpubJan-Mar (10.4161/pri.6.1.18272).

44. S. Keck, R. Nitsch, T. Grune, O. Ullrich, Proteasome inhibition by paired helical filament-tau in brains of patients with Alzheimer’s disease. Journal of neurochemistry 85, 115–122 (2003); published online EpubApr (

45. H. Snyder, K. Mensah, C. Theisler, J. Lee, A. Matouschek, B. Wolozin, Aggregated and monomeric alpha-synuclein bind to the S6’ proteasomal protein and inhibit proteasomal function. The Journal of biological chemistry 278, 11753–11759 (2003); published online EpubApr 04 (10.1074/jbc.M208641200).

46. Q. Guo, C. Lehmer, A. Martinez-Sanchez, T. Rudack, F. Beck, H. Hartmann, M. Perez-Berlanga, F. Frottin, M. S. Hipp, F. U. Hartl, D. Edbauer, W. Baumeister, R. Fernandez-Busnadiego, In Situ Structure of Neuronal C9orf72 Poly-GA Aggregates Reveals Proteasome Recruitment. Cell 172, 696–705 e612 (2018); published online EpubFeb 8 (10.1016/j.cell.2017.12.030).

47. T. A. Thibaudeau, R. T. Anderson, D. M. Smith, A common mechanism of proteasome impairment by neurodegenerative disease-associated oligomers. Nat Commun 9, 1097 (2018); published online EpubMar 15 (10.1038/s41467-018-03509-0).

48. A. W. Schaler, N. Myeku, Cilostazol, a phosphodiesterase 3 inhibitor, activates proteasome-mediated proteolysis and attenuates tauopathy and cognitive decline. Translational research : the journal of laboratory and clinical medicine 193, 31–41 (2018); published online EpubMar (10.1016/j.trsl.2017.11.004).

49. S. Lokireddy, N. V. Kukushkin, A. L. Goldberg, cAMP-induced phosphorylation of 26S proteasomes on Rpn6/PSMD11 enhances their activity and the degradation of misfolded proteins. Proceedings of the National Academy of Sciences of the United States of America 112, E7176–7185 (2015); published online EpubDec 29(10.1073/pnas.1522332112).

50. J. J. S. VerPlank, S. Lokireddy, J. Zhao, A. L. Goldberg, 26S Proteasomes are rapidly activated by diverse hormones and physiological states that raise cAMP and cause Rpn6 phosphorylation. Proc Natl Acad Sci U S A, (2019); published online EpubFeb 19 (10.1073/pnas.1809254116).

51. H. Zhang, B. Pan, P. Wu, N. Parajuli, M. D. Rekhter, A. L. Goldberg, X. Wang, PDE1 inhibition facilitates proteasomal degradation of misfolded proteins and protects against cardiac proteinopathy. Sci Adv 5, eaaw5870 (2019); published online EpubMay (10.1126/sciadv.aaw5870).

52. M. J. Ranek, E. J. Terpstra, J. Li, D. A. Kass, X. Wang, Protein kinase g positively regulates proteasome-mediated degradation of misfolded proteins. Circulation 128, 365–376 (2013); published online EpubJul 23 (10.1161/CIRCULATIONAHA.113.001971).

53. X. Guo, X. Wang, Z. Wang, S. Banerjee, J. Yang, L. Huang, J. E. Dixon, Site-specific proteasome phosphorylation controls cell proliferation and tumorigenesis. Nat Cell Biol 18, 202–212 (2016); published online EpubFeb (10.1038/ncb3289).

54. S. Banerjee, C. Ji, J. E. Mayfield, A. Goel, J. Xiao, J. E. Dixon, X. Guo, Ancient drug curcumin impedes 26S proteasome activity by direct inhibition of dual-specificity tyrosine-regulated kinase 2. Proc Natl Acad Sci U S A 115, 8155–8160 (2018); published online EpubAug 7 (10.1073/pnas.1806797115).

55. Y. Huang, A. Thathiah, Regulation of neuronal communication by G protein-coupled receptors. FEBS letters 589, 1607–1619 (2015); published online EpubJun 22 (10.1016/j.febslet.2015.05.007).

56. K. M. Betke, C. A. Wells, H. E. Hamm, GPCR mediated regulation of synaptic transmission. Progress in neurobiology 96, 304–321 (2012); published online EpubMar (10.1016/j.pneurobio.2012.01.009).

57. K. J. Gerber, K. E. Squires, J. R. Hepler, Roles for Regulator of G Protein Signaling Proteins in Synaptic Signaling and Plasticity. Molecular pharmacology 89, 273–286 (2016); published online EpubFeb (10.1124/mol.115.102210).

58. D. Vaudry, A. Falluel-Morel, S. Bourgault, M. Basille, D. Burel, O. Wurtz, A. Fournier, B. K. Chow, H. Hashimoto, L. Galas, H. Vaudry, Pituitary adenylate cyclase-activating polypeptide and its receptors: 20 years after the discovery. Pharmacol Rev 61, 283–357 (2009); published online EpubSep (10.1124/pr.109.001370).

59. T. Hirabayashi, T. Nakamachi, S. Shioda, Discovery of PACAP and its receptors in the brain. J Headache Pain 19, 28 (2018); published online EpubApr 4 (10.1186/s10194-018-0855-1).

60. S. Shioda, H. Ohtaki, T. Nakamachi, K. Dohi, J. Watanabe, S. Nakajo, S. Arata, S. Kitamura, H. Okuda, F. Takenoya, Y. Kitamura, Pleiotropic functions of PACAP in the CNS: neuroprotection and neurodevelopment. Ann N Y Acad Sci 1070, 550–560 (2006); published online EpubJul (10.1196/annals.1317.080).

61. S. Konzack, E. Thies, A. Marx, E. M. Mandelkow, E. Mandelkow, Swimming against the tide: mobility of the microtubule-associated protein tau in neurons. J Neurosci 27, 9916–9927 (2007); published online EpubSep 12 (10.1523/JNEUROSCI.0927-07.2007).

62. H. Zempel, F. J. A. Dennissen, Y. Kumar, J. Luedtke, J. Biernat, E. M. Mandelkow, E. Mandelkow, Axodendritic sorting and pathological missorting of Tau are isoform-specific and determined by axon initial segment architecture. J Biol Chem 292, 12192–12207 (2017); published online EpubJul 21 (10.1074/jbc.M117.784702).

63. A. Migheli, M. Butler, K. Brown, M. L. Shelanski, Light and electron microscope localization of the microtubule-associated tau protein in rat brain. J Neurosci 8, 1846–1851 (1988); published online EpubJun (

64. K. S. Kosik, A. Caceres, Tau protein and the establishment of an axonal morphology. J Cell Sci Suppl 15, 69–74 (1991).

65. S. L. DeVos, B. T. Corjuc, D. H. Oakley, C. K. Nobuhara, R. N. Bannon, A. Chase, C. Commins, J. A. Gonzalez, P. M. Dooley, M. P. Frosch, B. T. Hyman, Synaptic Tau Seeding Precedes Tau Pathology in Human Alzheimer’s Disease Brain. Frontiers in neuroscience 12, 267 (2018)10.3389/fnins.2018.00267).

66. H. Zempel, E. M. Mandelkow, Tau missorting and spastin-induced microtubule disruption in neurodegeneration: Alzheimer Disease and Hereditary Spastic Paraplegia. Molecular neurodegeneration 10, 68 (2015); published online EpubDec 21 (10.1186/s13024-015-0064-1).

67. E. Thies, E. M. Mandelkow, Missorting of tau in neurons causes degeneration of synapses that can be rescued by the kinase MARK2/Par-1. The Journal of neuroscience : the official journal of the Society for Neuroscience 27, 2896–2907 (2007); published online EpubMar 14 (10.1523/JNEUROSCI.4674-06.2007).

68. B. Dejanovic, M. A. Huntley, A. De Maziere, W. J. Meilandt, T. Wu, K. Srinivasan, Z. Jiang, V. Gandham, B. A. Friedman, H. Ngu, O. Foreman, R. A. D. Carano, B. Chih, J. Klumperman, C. Bakalarski, J. E. Hanson, M. Sheng, Changes in the Synaptic Proteome in Tauopathy and Rescue of Tau-Induced Synapse Loss by C1q Antibodies. Neuron 100, 1322–1336 e1327 (2018); published online EpubDec 19 (10.1016/j.neuron.2018.10.014).

69. K. Eckermann, M. M. Mocanu, I. Khlistunova, J. Biernat, A. Nissen, A. Hofmann, K. Schonig, H. Bujard, A. Haemisch, E. Mandelkow, L. Zhou, G. Rune, E. M. Mandelkow, The beta-propensity of Tau determines aggregation and synaptic loss in inducible mouse models of tauopathy. J Biol Chem 282, 31755–31765 (2007); published online EpubOct 26 (10.1074/jbc.M705282200).

70. J. S. Jackson, J. Witton, J. D. Johnson, Z. Ahmed, M. Ward, A. D. Randall, M. L. Hutton, J. T. Isaac, M. J. O’Neill, M. C. Ashby, Altered Synapse Stability in the Early Stages of Tauopathy. Cell reports 18, 3063–3068 (2017); published online EpubMar 28 (10.1016/j.celrep.2017.03.013).

71. L. Zhou, J. McInnes, K. Wierda, M. Holt, A. G. Herrmann, R. J. Jackson, Y. C. Wang, J. Swerts, J. Beyens, K. Miskiewicz, S. Vilain, I. Dewachter, D. Moechars, B. De Strooper, T. L. Spires-Jones, J. De Wit, P. Verstreken, Tau association with synaptic vesicles causes presynaptic dysfunction. Nat Commun 8, 15295 (2017); published online EpubMay 11 (10.1038/ncomms15295).

72. N. A. Hoffmann, M. M. Dorostkar, S. Blumenstock, M. Goedert, J. Herms, Impaired plasticity of cortical dendritic spines in P301S tau transgenic mice. Acta Neuropathol Commun 1, 82 (2013); published online EpubDec 17 (10.1186/2051-5960-1-82).

73. Y. Yoshiyama, M. Higuchi, B. Zhang, S. M. Huang, N. Iwata, T. C. Saido, J. Maeda, T. Suhara, J. Q. Trojanowski, V. M. Lee, Synapse loss and microglial activation precede tangles in a P301S tauopathy mouse model. Neuron 53, 337–351 (2007); published online EpubFeb 1 (10.1016/j.neuron.2007.01.010).

74. L. M. Ittner, Y. D. Ke, F. Delerue, M. Bi, A. Gladbach, J. van Eersel, H. Wolfing, B. C. Chieng, M. J. Christie, I. A. Napier, A. Eckert, M. Staufenbiel, E. Hardeman, J. Gotz, Dendritic function of tau mediates amyloid-beta toxicity in Alzheimer’s disease mouse models. Cell 142, 387–397 (2010); published online EpubAug 6 (10.1016/j.cell.2010.06.036).

75. M. A. Chabrier, D. Cheng, N. A. Castello, K. N. Green, F. M. LaFerla, Synergistic effects of amyloid-beta and wild-type human tau on dendritic spine loss in a floxed double transgenic model of Alzheimer’s disease. Neurobiology of disease 64, 107–117 (2014); published online EpubApr (10.1016/j.nbd.2014.01.007).

76. A. Van der Jeugd, K. Hochgrafe, T. Ahmed, J. M. Decker, A. Sydow, A. Hofmann, D. Wu, L. Messing, D. Balschun, R. D’Hooge, E. M. Mandelkow, Cognitive defects are reversible in inducible mice expressing pro-aggregant full-length human Tau. Acta neuropathologica 123, 787–805 (2012); published online EpubJun (10.1007/s00401-012-0987-3).

77. H. Braak, K. Del Tredici, The preclinical phase of the pathological process underlying sporadic Alzheimer’s disease. Brain : a journal of neurology 138, 2814–2833 (2015); published online EpubOct (10.1093/brain/awv236).

78. L. I. Binder, A. L. Guillozet-Bongaarts, F. Garcia-Sierra, R. W. Berry, Tau, tangles, and Alzheimer’s disease. Biochimica et biophysica acta 1739, 216–223 (2005); published online EpubJan 3 (10.1016/j.bbadis.2004.08.014).

79. S. Takeda, S. Wegmann, H. Cho, S. L. DeVos, C. Commins, A. D. Roe, S. B. Nicholls, G. A. Carlson, R. Pitstick, C. K. Nobuhara, I. Costantino, M. P. Frosch, D. J. Muller, D. Irimia, B. T. Hyman, Neuronal uptake and propagation of a rare phosphorylated high-molecular-weight tau derived from Alzheimer’s disease brain. Nature communications 6, 8490 (2015); published online EpubOct 13 (10.1038/ncomms9490).

80. M. C. de Wilde, C. R. Overk, J. W. Sijben, E. Masliah, Meta-analysis of synaptic pathology in Alzheimer’s disease reveals selective molecular vesicular machinery vulnerability. Alzheimer’s & dementia : the journal of the Alzheimer’s Association 12, 633–644 (2016); published online EpubJun (10.1016/j.jalz.2015.12.005).

81. J. Brettschneider, K. Del Tredici, V. M. Lee, J. Q. Trojanowski, Spreading of pathology in neurodegenerative diseases: a focus on human studies. Nature reviews. Neuroscience 16, 109–120 (2015); published online EpubFeb (10.1038/nrn3887).

82. T. L. Spires-Jones, B. T. Hyman, The intersection of amyloid beta and tau at synapses in Alzheimer’s disease. Neuron 82, 756–771 (2014); published online EpubMay 21 (10.1016/j.neuron.2014.05.004).

83. J. L. Furman, J. Vaquer-Alicea, C. L. White, 3rd, N. J. Cairns, P. T. Nelson, M. I. Diamond, Widespread tau seeding activity at early Braak stages. Acta neuropathologica 133, 91–100 (2017); published online EpubJan (10.1007/s00401-016-1644-z).

84. J. Vaquer-Alicea, M. I. Diamond, Propagation of Protein Aggregation in Neurodegenerative Diseases. Annu Rev Biochem 88, 785–810 (2019); published online EpubJun 20 (10.1146/annurev-biochem-061516-045049).

85. J. W. Wu, M. Herman, L. Liu, S. Simoes, C. M. Acker, H. Figueroa, J. I. Steinberg, M. Margittai, R. Kayed, C. Zurzolo, G. Di Paolo, K. E. Duff, Small misfolded Tau species are internalized via bulk endocytosis and anterogradely and retrogradely transported in neurons. The Journal of biological chemistry 288, 1856–1870 (2013); published online EpubJan 18 (10.1074/jbc.M112.394528).

86. A. Mercer, H. Ronnholm, J. Holmberg, H. Lundh, J. Heidrich, O. Zachrisson, A. Ossoinak, J. Frisen, C. Patrone, PACAP promotes neural stem cell proliferation in adult mouse brain. J Neurosci Res 76, 205–215 (2004); published online EpubApr 15 (10.1002/jnr.20038).

87. S. Ohta, C. Gregg, S. Weiss, Pituitary adenylate cyclase-activating polypeptide regulates forebrain neural stem cells and neurogenesis in vitro and in vivo. J Neurosci Res 84, 1177–1186 (2006); published online EpubNov 1 (10.1002/jnr.21026).

88. T. Seaborn, O. Masmoudi-Kouli, A. Fournier, H. Vaudry, D. Vaudry, Protective effects of pituitary adenylate cyclase-activating polypeptide (PACAP) against apoptosis. Curr Pharm Des 17, 204–214 (2011).

89. D. Vaudry, C. Rousselle, M. Basille, A. Falluel-Morel, T. F. Pamantung, M. Fontaine, A. Fournier, H. Vaudry, B. J. Gonzalez, Pituitary adenylate cyclase-activating polypeptide protects rat cerebellar granule neurons against ethanol-induced apoptotic cell death. Proc Natl Acad Sci U S A 99, 6398–6403 (2002); published online EpubApr 30 (10.1073/pnas.082112699).

90. K. Szabadfi, A. Szabo, P. Kiss, D. Reglodi, G. Setalo, Jr., K. Kovacs, A. Tamas, G. Toth, R. Gabriel, PACAP promotes neuron survival in early experimental diabetic retinopathy. Neurochem Int 64, 84–91 (2014); published online EpubJan (10.1016/j.neuint.2013.11.005).

91. K. Hajji, A. Mteyrek, J. Sun, M. Cassar, S. Mezghani, J. Leprince, D. Vaudry, O. Masmoudi-Kouki, S. Birman, Neuroprotective effects of PACAP against paraquat-induced oxidative stress in the Drosophila central nervous system. Hum Mol Genet 28, 1905–1918 (2019); published online EpubJun 1 (10.1093/hmg/ddz031).

92. H. Ohtaki, T. Nakamachi, K. Dohi, Y. Aizawa, A. Takaki, K. Hodoyama, S. Yofu, H. Hashimoto, N. Shintani, A. Baba, M. Kopf, Y. Iwakura, K. Matsuda, A. Arimura, S. Shioda, Pituitary adenylate cyclase-activating polypeptide (PACAP) decreases ischemic neuronal cell death in association with IL-6. Proc Natl Acad Sci U S A 103, 7488–7493 (2006); published online EpubMay 9 (10.1073/pnas.0600375103).

93. S. K. Kaufman, D. W. Sanders, T. L. Thomas, A. J. Ruchinskas, J. Vaquer-Alicea, A. M. Sharma, T. M. Miller, M. I. Diamond, Tau Prion Strains Dictate Patterns of Cell Pathology, Progression Rate, and Regional Vulnerability In Vivo. Neuron 92, 796–812 (2016); published online EpubNov 23 (10.1016/j.neuron.2016.09.055).

94. P. Robberecht, P. Gourlet, P. De Neef, M. C. Woussen-Colle, M. C. Vandermeers-Piret, A. Vandermeers, J. Christophe, Structural requirements for the occupancy of pituitary adenylate-cyclase-activating-peptide (PACAP) receptors and adenylate cyclase activation in human neuroblastoma NB-OK-1 cell membranes. Discovery of PACAP(6-38) as a potent antagonist. European journal of biochemistry 207, 239–246 (1992); published online EpubJul 1 (10.1111/j.1432-1033.1992.tb17043.x).

95. A. Arimura, A. Somogyvari-Vigh, A. Miyata, K. Mizuno, D. H. Coy, C. Kitada, Tissue distribution of PACAP as determined by RIA: highly abundant in the rat brain and testes. Endocrinology 129, 2787–2789 (1991); published online EpubNov (10.1210/endo-129-5-2787).

96. M. A. Ghatei, K. Takahashi, Y. Suzuki, J. Gardiner, P. M. Jones, S. R. Bloom, Distribution, molecular characterization of pituitary adenylate cyclase-activating polypeptide and its precursor encoding messenger RNA in human and rat tissues. J Endocrinol 136, 159–166 (1993); published online EpubJan (10.1677/joe.0.1360159).

97. H. Hashimoto, H. Nogi, K. Mori, H. Ohishi, R. Shigemoto, K. Yamamoto, T. Matsuda, N. Mizuno, S. Nagata, A. Baba, Distribution of the mRNA for a pituitary adenylate cyclase-activating polypeptide receptor in the rat brain: an in situ hybridization study. J Comp Neurol 371, 567–577 (1996); published online EpubAug 5 (10.1002/(SICI)1096-9861(19960805)371:4<567::AID-CNE6>3.0.CO;2-2).

98. M. Nomura, Y. Ueta, R. Serino, N. Kabashima, I. Shibuya, H. Yamashita, PACAP type I receptor gene expression in the paraventricular and supraoptic nuclei of rats. Neuroreport 8, 67–70 (1996); published online EpubDec 20 (10.1097/00001756-199612200-00014).

99. S. Shioda, Y. Nakai, S. Nakajo, K. Nakaya, A. Arimura, Localization of pituitary adenylate cyclase-activating polypeptide and its type I receptors in the rat ovary: immunohistochemistry and in situ hybridization. Ann N Y Acad Sci 805, 677–683 (1996); published online EpubDec 26 (10.1111/j.1749-6632.1996.tb17540.x).

100. S. Shioda, Y. Shuto, A. Somogyvari-Vigh, G. Legradi, H. Onda, D. H. Coy, S. Nakajo, A. Arimura, Localization and gene expression of the receptor for pituitary adenylate cyclase-activating polypeptide in the rat brain. Neuroscience research 28, 345–354 (1997); published online EpubAug (

101. M. Basille, D. Cartier, D. Vaudry, I. Lihrmann, A. Fournier, P. Freger, N. Gallo-Payet, H. Vaudry, B. Gonzalez, Localization and characterization of pituitary adenylate cyclase-activating polypeptide receptors in the human cerebellum during development. J Comp Neurol 496, 468–478 (2006); published online EpubJun 1 (10.1002/cne.20934).

102. P. Han, R. J. Caselli, L. Baxter, G. Serrano, J. Yin, T. G. Beach, E. M. Reiman, J. Shi, Association of pituitary adenylate cyclase-activating polypeptide with cognitive decline in mild cognitive impairment due to Alzheimer disease. JAMA Neurol 72, 333–339 (2015); published online EpubMar (10.1001/jamaneurol.2014.3625).

103. P. Han, Z. Tang, J. Yin, M. Maalouf, T. G. Beach, E. M. Reiman, J. Shi, Pituitary adenylate cyclase-activating polypeptide protects against beta-amyloid toxicity. Neurobiology of aging 35, 2064–2071 (2014); published online EpubSep (10.1016/j.neurobiolaging.2014.03.022).

104. A. Thathiah, B. De Strooper, The role of G protein-coupled receptors in the pathology of Alzheimer’s disease. Nature reviews. Neuroscience 12, 73–87 (2011); published online EpubFeb (10.1038/nrn2977).

105. J. R. Fisher, C. E. Wallace, D. L. Tripoli, Y. I. Sheline, J. R. Cirrito, Redundant Gs-coupled serotonin receptors regulate amyloid-beta metabolism in vivo. Molecular neurodegeneration 11, 45 (2016); published online EpubJun 18 (10.1186/s13024-016-0112-5).

106. N. Myeku, H. Wang, M. E. Figueiredo-Pereira, cAMP stimulates the ubiquitin/proteasome pathway in rat spinal cord neurons. Neuroscience letters 527, 126–131 (2012); published online EpubOct 11 (10.1016/j.neulet.2012.08.051).

107. N. V. K. Sudarsanareddy Lokireddy, Alfred L. Goldberg, cAMP-induced phosphorylation of 26S proteasomes on Rpn6/PSMD11 enhances their activity and the degradation of misfolded proteins. Proc. Natl. Acad. Sci, (2015) 10.1073/pnas.1522332112).

108. L. A. Merriam, C. N. Baran, B. M. Girard, J. C. Hardwick, V. May, R. L. Parsons, Pituitary adenylate cyclase 1 receptor internalization and endosomal signaling mediate the pituitary adenylate cyclase activating polypeptide-induced increase in guinea pig cardiac neuron excitability. J Neurosci 33, 4614–4622 (2013); published online EpubMar 6 (10.1523/JNEUROSCI.4999-12.2013).

109. J. D. Tompkins, T. A. Clason, T. R. Buttolph, B. M. Girard, A. K. Linden, J. C. Hardwick, L. A. Merriam, V. May, R. L. Parsons, Src family kinase inhibitors blunt PACAP-induced PAC1 receptor endocytosis, phosphorylation of ERK, and the increase in cardiac neuron excitability. Am J Physiol Cell Physiol 314, C233–C241 (2018); published online EpubFeb 1 (10.1152/ajpcell.00223.2017).

110. V. May, T. R. Buttolph, B. M. Girard, T. A. Clason, R. L. Parsons, PACAP-induced ERK activation in HEK cells expressing PAC1 receptors involves both receptor internalization and PKC signaling. American journal of physiology. Cell physiology 306, C1068–1079 (2014); published online EpubJun 1 (10.1152/ajpcell.00001.2014).

111. H. C. Tai, H. Besche, A. L. Goldberg, E. M. Schuman, Characterization of the Brain 26S Proteasome and its Interacting Proteins. Front Mol Neurosci 3, (2010)10.3389/fnmol.2010.00012).

112. H. C. Besche, A. L. Goldberg, Affinity purification of mammalian 26S proteasomes using an ubiquitin-like domain. Methods in molecular biology 832, 423–432 (2012)10.1007/978-1-61779-474-2_29).

113. M. J. Polanco, S. Parodi, D. Piol, C. Stack, M. Chivet, A. Contestabile, H. C. Miranda, P. M. Lievens, S. Espinoza, T. Jochum, A. Rocchi, C. Grunseich, R. R. Gainetdinov, A. C. Cato, A. P. Lieberman, A. R. La Spada, F. Sambataro, K. H. Fischbeck, I. Gozes, M. Pennuto, Adenylyl cyclase activating polypeptide reduces phosphorylation and toxicity of the polyglutamine-expanded androgen receptor in spinobulbar muscular atrophy. Sci Transl Med 8, 370ra181 (2016); published online EpubDec 21 (10.1126/scitranslmed.aaf9526).

114. L. Blazquez-Llorca, V. Garcia-Marin, P. Merino-Serrais, J. Avila, J. DeFelipe, Abnormal tau phosphorylation in the thorny excrescences of CA3 hippocampal neurons in patients with Alzheimer’s disease. J Alzheimers Dis 26, 683–698 (2011)10.3233/JAD-2011-110659).

115. P. Merino-Serrais, R. Benavides-Piccione, L. Blazquez-Llorca, A. Kastanauskaite, A. Rabano, J. Avila, J. DeFelipe, The influence of phospho-tau on dendritic spines of cortical pyramidal neurons in patients with Alzheimer’s disease. Brain 136, 1913–1928 (2013); published online EpubJun (10.1093/brain/awt088).

116. N. O. Dalby, C. Volbracht, L. Helboe, P. H. Larsen, H. S. Jensen, J. Egebjerg, A. B. Elvang, Altered function of hippocampal CA1 pyramidal neurons in the rTg4510 mouse model of tauopathy. J Alzheimers Dis 40, 429–442 (2014)10.3233/JAD-131358).

117. D. L. Dickstein, H. Brautigam, S. D. Stockton, Jr., J. Schmeidler, P. R. Hof, Changes in dendritic complexity and spine morphology in transgenic mice expressing human wild-type tau. Brain Struct Funct 214, 161–179 (2010); published online EpubMar (10.1007/s00429-010-0245-1).

118. A. S. Hauser, M. M. Attwood, M. Rask-Andersen, H. B. Schioth, D. E. Gloriam, Trends in GPCR drug discovery: new agents, targets and indications. Nature reviews. Drug discovery, (2017); published online EpubOct 27 (10.1038/nrd.2017.178).

119. Y. Huang, N. Todd, A. Thathiah, The role of GPCRs in neurodegenerative diseases: avenues for therapeutic intervention. Current opinion in pharmacology 32, 96–110 (2017); published online EpubFeb (10.1016/j.coph.2017.02.001).

120. J. T. Lin, W. C. Chang, H. M. Chen, H. L. Lai, C. Y. Chen, M. H. Tao, Y. Chern, Regulation of feedback between protein kinase A and the proteasome system worsens Huntington’s disease. Molecular and cellular biology 33, 1073–1084 (2013); published online EpubMar (10.1128/MCB.01434-12).

121. F. J. Dennissen, M. Anglada-Huguet, A. Sydow, E. Mandelkow, E. M. Mandelkow, Adenosine A1 receptor antagonist rolofylline alleviates axonopathy caused by human Tau DeltaK280. Proc Natl Acad Sci U S A 113, 11597–11602 (2016); published online EpubOct 11 (10.1073/pnas.1603119113).

122. D. L. Smith, J. Pozueta, B. Gong, O. Arancio, M. Shelanski, Reversal of long-term dendritic spine alterations in Alzheimer disease models. Proc Natl Acad Sci U S A 106, 16877–16882 (2009); published online EpubSep 29 (10.1073/pnas.0908706106).

123. B. D. Green, N. Irwin, P. R. Flatt, Pituitary adenylate cyclase-activating peptide (PACAP): assessment of dipeptidyl peptidase IV degradation, insulin-releasing activity and antidiabetic potential. Peptides 27, 1349–1358 (2006); published online EpubJun (10.1016/j.peptides.2005.11.010).

124. M. Li, J. L. Maderdrut, J. J. Lertora, V. Batuman, Intravenous infusion of pituitary adenylate cyclase-activating polypeptide (PACAP) in a patient with multiple myeloma and myeloma kidney: a case study. Peptides 28, 1891–1895 (2007); published online EpubSep (10.1016/j.peptides.2007.05.002).

125. R. Morris, Developments of a water-maze procedure for studying spatial learning in the rat. Journal of neuroscience methods 11, 47–60 (1984); published online EpubMay (

126. S. L. DeVos, T. M. Miller, Direct intraventricular delivery of drugs to the rodent central nervous system. Journal of visualized experiments : JoVE, e50326 (2013); published online EpubMay 12 (10.3791/50326).

127. C. M. Acker, S. K. Forest, R. Zinkowski, P. Davies, C. d’Abramo, Sensitive quantitative assays for tau and phospho-tau in transgenic mouse models. Neurobiology of aging 34, 338–350 (2013); published online EpubJan (10.1016/j.neurobiolaging.2012.05.010).

